# Activity-dependent expression of *Fezf2* regulates inhibitory synapse formation in pyramidal cells

**DOI:** 10.1101/2024.01.11.575171

**Authors:** Tim Kroon, Stella Sanalidou, Patricia Maeso, Teresa Garces, Han Yang, Beatriz Rico

## Abstract

The function of the cerebral cortex relies on the precise integration of diverse neuronal populations during development, which is regulated by dynamic fine-tuning mechanisms maintaining the balance between excitation and inhibition. For instance, the development of excitatory pyramidal cells is simultaneously and precisely counterbalanced by the formation of inhibitory synapses during the maturation of neuronal circuits. Although this process relies on neuronal activity, different types of pyramidal cells likely respond to changes in activity through the expression of cell-specific genes. However, the molecular programs underlying the activity-dependent recruitment of inhibition by distinct types of pyramidal cells in the neocortex are unknown. Here, we combined neuronal activity manipulation with ribosome-associated mRNA profiling of layer 5 (L5) extra-telencephalic (ET) cells to address this question in mice. We unveiled a novel function for the selector gene *Fezf2* as an activity-dependent transcription factor controlling the parvalbumin inputs onto L5 ET neurons. One of the downstream effectors of FEZF2 shaping the formation of inhibitory synapses onto L5 ET pyramidal cells is the cell-surface molecule cadherin 22. Our study identifies activity-dependent factors regulating the cell type-specific assembly of inhibitory synapses onto pyramidal cells.

## Introduction

The function of the cerebral cortex relies on the precise interplay between two neuronal populations: excitatory glutamatergic principal or pyramidal cells and inhibitory gamma-aminobutyric acid-expressing (GABAergic) interneurons. The connectivity motifs established between these two neuronal populations are highly specialized and crucial for the function of cortical circuits. For example, different types of inhibitory interneurons have been shown to target specific subcellular compartments of principal cells, increasing the computational power of cortical circuitries^1–3^.

A remarkable diversity of molecular programs supports the exquisite variety of cortical inhibitory connectivity that emerges during development. The prevailing view conveys that synaptic development has two components: (1) a genetically determined process sustained by specific molecular codes and (2) an activity-dependent mechanism that shapes the circuitries ^4^. Several cell-surface molecule complexes are required for the development of GABAergic synapses, including Neuroligin-2, Sema4D-PlexinB1, Nrg1/3-ErbB4, Slitrk3-PTPδ and Neurexin2αß-IgsF21^5–10^. Moreover, the specific cellular and subcellular targeting of pyramidal cells by interneurons is also regulated by cell-surface molecules^11, 12^. The development of inhibitory synapses in the cerebral cortex unfolds through a protracted period of postnatal development^11, 13, 14^ and activity sculpts the inhibitory cortical circuitries until they reach the mature state^13, 15, 16^.

In the adult brain, activity-dependent mechanisms are fundamental to maintaining reliable coding during fluctuating activity patterns, such as those caused by sensory experience ^17^. Pyramidal neurons also regulate the inhibition they receive through homeostatic mechanisms^18–21^. Synaptic and cell-intrinsic changes underlying homeostatic plasticity are mediated through the activity-dependent expression of immediate early genes (IEGs). These transcription factors promote the subsequent expression of late response genes (LRGs), which often encode secreted molecules or proteins with synaptic localization^22^. An emerging view is that while IEGs are relatively common across different neuronal classes, LRGs induced in response to activity are cell type-specific^23^. Yet, how broadly expressed IEGs regulate cell type-specific LRGs and to what extent the mechanisms regulating activity-dependent gene expression during postnatal development are like those described in the adult brain remains to be elucidated.

Cortical inhibitory synapses onto pyramidal cells undergo a steady increase during the first month of development^11, 14^, probably matching the progressive surge in excitatory inputs and activity among pyramidal cells^24, 25^. Consistently, in vitro experiments revealed that the formation of inhibitory synapses onto excitatory neurons is regulated by activity^13, 26^, and in the hippocampus, it requires *Npas4*, an IEG^26^. However, despite the relevance of inhibitory synapses and activity in the maturation of cortical circuitries^15^, how pyramidal cells respond to the substantial increase in activity they receive during development and the molecular programs governing these processes are still poorly understood.

Here, we investigated these questions by focusing on L5 ET pyramidal cells, the most excitable cortical pyramidal neurons^27–29^. L5 ET pyramidal cells receive remarkably more PV+ basket inhibitory inputs than other pyramidal cells since early postnatal development^12^. Combining manipulation of neuronal activity with ribosome-associated mRNA profiling, we identified a novel role for selector gene *Fezf2* as an activity-dependent transcription factor regulating the inhibitory wiring of L5 ET neurons. We also revealed that the expression of *Fezf2* is not triggered by activity in other pyramidal cells. In response to activity, FEZF2 regulates PV+ perisomatic inhibition onto L5 ET pyramidal cells without affecting other features of cell identity. Moreover, we uncovered that the cell-adhesion molecule Cadherin-22 (*Cdh22*) functions downstream of *Fezf2* and regulates the density of PV+ inhibitory inputs onto ET pyramidal cells. Altogether, our findings unveil a novel function for FEZF2 as an activity-dependent transcription factor that governs inhibitory synapses onto pyramidal cells in a cell type-specific manner.

## Results

### Neuronal activity regulates the development of inhibitory inputs onto pyramidal cells

To investigate whether activity regulates the formation of PV+ perisomatic inhibitory synapses, we focused on L5 ET pyramidal cells. These neurons receive a high density of perisomatic inhibition that increases steadily during the first month of postnatal life in mice^12^. We sought to reduce the excitability of L5 ET pyramidal cells during early postnatal development by expressing the inwardly rectifying potassium channel Kir2.1^30^. We used an intersectional strategy to label L5 ET pyramidal cells in the primary somatosensory cortex (S1). In brief, we simultaneously injected a retrograde adeno-associated virus (AAV) expressing Cre recombinase (*rAAV_retro_-Cre*) in the pontine nuclei of postnatal day (P) 1 wild-type mice and a Cre-dependent AAV expressing Kir2.1 and tdTomato in the S1 ipsilateral to the pons injection (Fig.1a and 1b). We used this approach in all subsequent experiments to label L5 ET neurons unless stated otherwise.

To confirm that Kir2.1 expression lowers the excitability of L5 ET cells before the initial surge of inhibitory synapse formation around P10^11^, we prepared acute slices from Kir2.1-expressing mice at P8 and performed electrophysiological recordings of infected and neighboring uninfected deep L5 ET pyramidal cells. Uninfected L5 ET cells were identified by their large soma and pronounced sag current, characteristic of these neurons^31^. Compared to uninfected cells, the resting membrane potential (RMP) of Kir2.1-expressing cells was hyperpolarized, and the input resistance (Ri) decreased, leading to a marked increase in rheobase that denoted a decrease in excitability (Fig. 1c and Extended Data Fig. 1a, b).

**Fig. 1:**
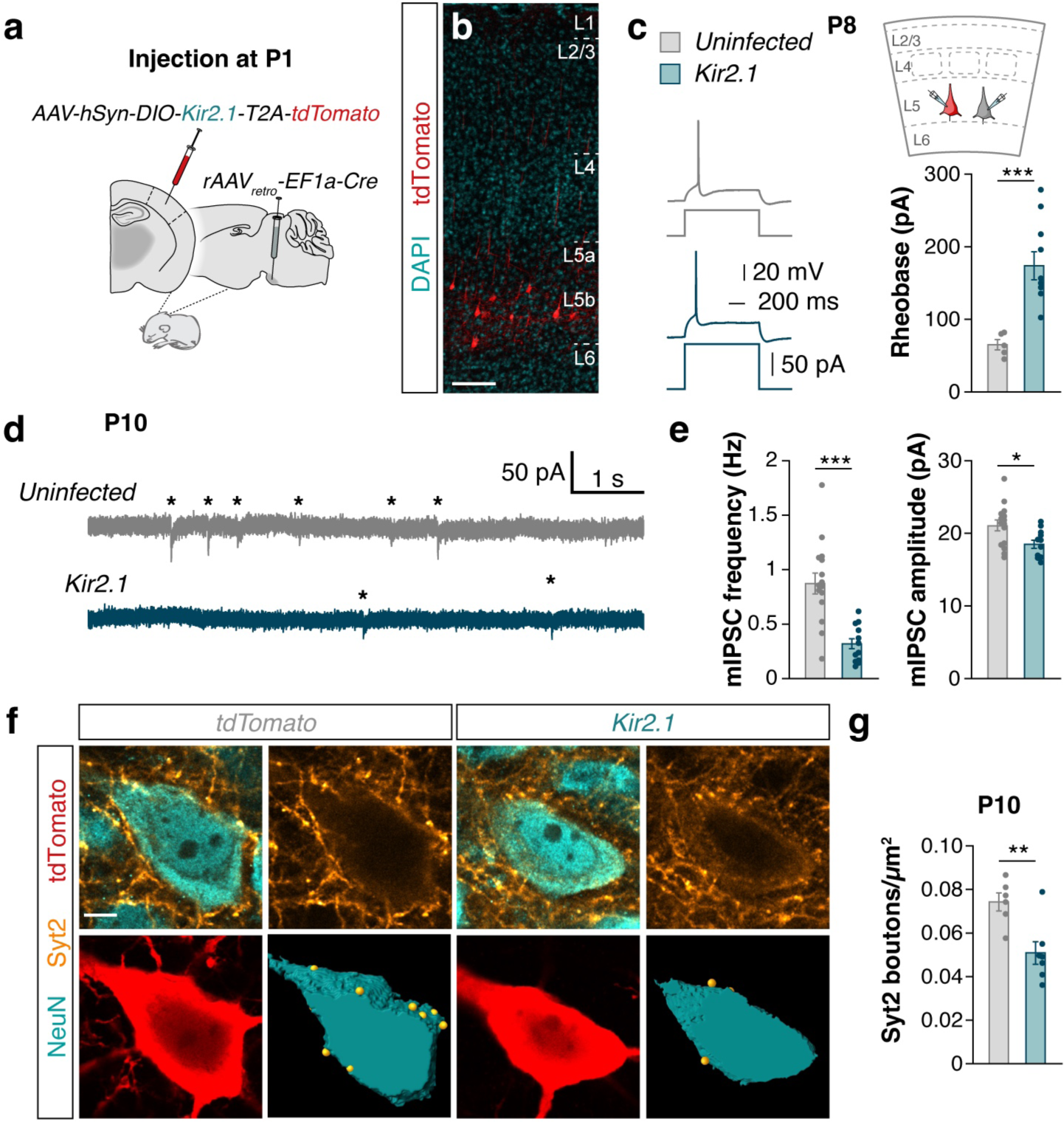
Decreased neuronal activity reduces inhibitory inputs during development. **a**, Experimental design for decreasing activity with Kir2.1. **b**, Expression of tdTomato in AAV-infected cells in deep L5. **c**, Electrophysiological measurement of excitability of infected vs uninfected ET pyramidal cells. Left, example traces of the first current injection to elicit an action potential. Bottom right, quantification of rheobase (uninfected, *n* = 5 cells from 2 animals; Kir2.1 *n* = 10 cells from 2 animals, two-tailed Student’s *t*-test, \*\*\**P* < 0.001). See also Fig. S1. **d**, Example traces of mIPSCs recorded from uninfected and Kir2.1-expressing cells. Asterisks indicate the onset of mIPSC events. **e**, Quantification of mIPSC frequency (uninfected, *n* = 16 cells from 6 animals; Kir2.1, *n* = 13 cells from 6 animals, two-tailed Student’s *t*-test, \*\*\**P* < 0.001) and amplitude (uninfected, *n* = 15 cells from 6 animals; Kir2.1, *n* = 12 cells from 6 animals, two-tailed Student’s *t*-test, \**P* < 0.05). **f**, Single-plane confocal images showing presynaptic Syt2 puncta (yellow), NeuN (cyan) and tdTomato (red). Bottom right: 3-dimensional Imaris reconstruction of the soma and Syt2 puncta. **g**, Quantification of the density of Syt2 puncta onto the soma (tdTomato, *n* = 6 animals; Kir2.1, *n* = 7 animals, two-tailed Student’s *t*-test, \*\**P* < 0.01). Data are mean ± SEM. Scale bars, 100 μm (**b**), 5 μm (**f**).

To establish whether the observed reduction in excitability leads to a functional decrease in inhibitory inputs, we measured miniature inhibitory postsynaptic currents (mIPSCs). We found a reduction in the frequency and amplitude of mIPSCs in Kir2.1-expressing cells compared to uninfected neighboring L5 ET pyramidal cells at P10 (Fig. 1d, e). Control experiments revealed that expression of a mutated version of Kir2.1 (mutKir2.1) encoding a non-conducting channel^19^ did not reduce the frequency or amplitude of mIPSCs compared to neighboring uninfected cells (Extended Data Fig. 1c-e). We next asked whether the decrease in inhibition was due to a reduction in the number of PV+ inputs, the primary source of perisomatic inhibition of these neurons^12^. To this end, we labeled control and Kir2.1-expressing L5 ET cells with tdTomato and quantified the density of perisomatic PV+ boutons using the presynaptic marker synaptotagmin-2 (Syt2), which is specifically expressed in PV+ basket cells^32^. We observed a reduction in the density of Syt2+ boutons contacting the soma of L5 ET neurons expressing Kir2.1 compared to control neurons at P10 (Fig. 1f, g). Altogether, these experiments revealed that decreasing the activity of L5 ET pyramidal cells during early postnatal development reduces the number of PV+ basket cell synapses these cells receive, demonstrating that neuronal activity regulates inhibitory synapse development in these neurons.

### *Fezf2* is regulated by neuronal activity in a cell type-specific manner

To identify the activity-dependent molecular mechanisms regulating the development of PV+ perisomatic inhibition onto L5 ET pyramidal neurons, we analyzed changes in the translatome of L5 ET neurons following the modulation of their activity during postnatal development. To this end, we used viral translating ribosome affinity purification (vTRAP)^33^ to isolate ribosome-associated mRNA from pontine-projecting ET neurons expressing Kir2.1 or mutKir2.1. In brief, we generated Cre-dependent AAVs expressing hemagglutinin (HA)-tagged ribosomal protein (Rpl10a) and either Kir2.1 (Kir2.1-vTRAP) or mutKir2.1 (mutKir2.1-vTRAP) (Extended Data Fig2a). To verify that Kir2.1-vTRAP decreases the excitability of L5 ET cells, we injected Kir2.1-vTRAP and mutKir2.1-vTRAP AAVs in *Rbp4^Cre/+^* mice^34^, in which Cre is expressed in most L5 pyramidal cells (Extended Data Fig. 2b). We then prepared acute slices and filled recorded cells with neurobiotin to verify post-hoc that they expressed the corresponding virus using immunohistochemistry against the HA-tag (Extended Data Fig2c). As expected, the excitability of cells expressing Kir2.1-vTRAP and the density of PV+ basket cell inputs they received were decreased compared to neurons expressing mutKir2.1-vTRAP (Extended Data Fig2d-h).

We isolated ribosome-associated mRNAs from L5 ET pyramidal cells expressing the Kir2.1-vTRAP or mutKir2.1-vTRAP at P10 and performed RNA-sequencing (Fig. 2a and Extended Data Fig. 3). We identified 673 differentially expressed genes (DEGs) between control and Kir2.1-expressing ET neurons (Fig. 2b and Extended Data Fig. 3). Gene ontology (GO) analysis of DEGs revealed enrichment in synaptic processes (Fig. 2c). We focused on transcription factors and genes directly involved in synaptic assembly and synaptic maintenance by selecting specific Panther GO terms (see Methods). We then ranked the selected genes using three criteria: (1) fold-change differential expression between Kir2.1- and mutKir2.1-expressing neurons; (2) gene expression levels; and (3) enrichment in ET cells compared to IT cells (Fig. 2d, see Methods for details). Top-ranking genes from this analysis include several cell-surface adhesion molecules and the zinc-finger transcription factor *Fezf2*. This selector gene is well known for its involvement in the early specification of L5 ET pyramidal cells^35–39^, so we were intrigued by the observation that the expression of *Fezf2* might be controlled in an activity-dependent manner in the postnatal cortex.

**Fig. 2:**
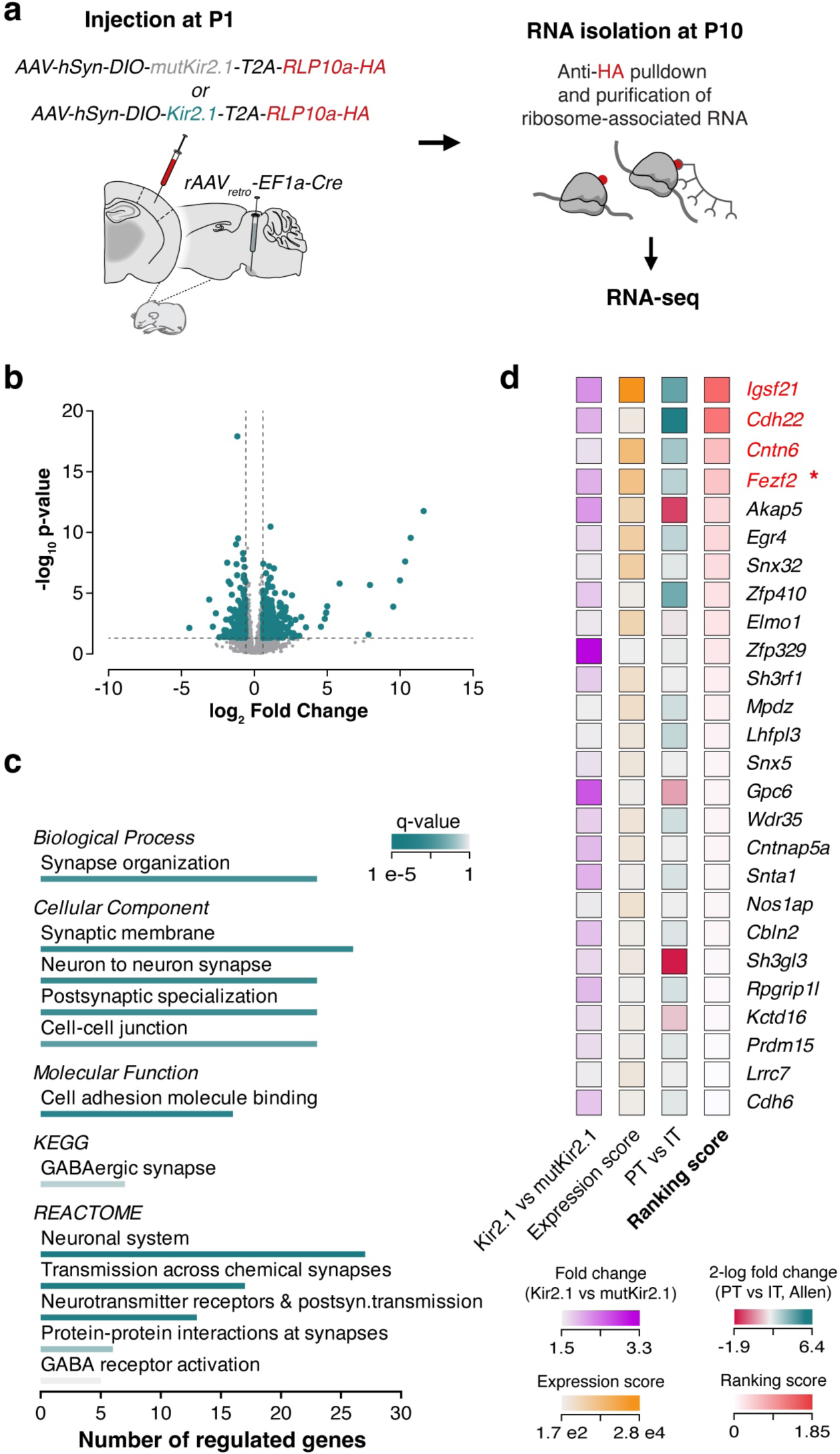
Transcriptional programs regulated by activity in L5 ET pyramidal cells. **a**, Experimental design for the profiling of ribosome-associated mRNA. **b**, Volcano plot displaying significantly differentially expressed ribosome-associated RNAs (teal) in ET pyramidal cells expressing Kir2.1 compared to mutKir2.1 controls. Each dot represents one gene. Criteria: fold change > 1.5, p-value < 0.05. **c**, Selected Gene Ontology (GO) terms significantly enriched in the dataset of differentially regulated genes in ET neurons expressing Kir2.1 compared to mutKir2.1 controls. Headers indicate GO domain/database. Color indicates the q-value, and the length of the bar indicates the number of regulated genes within the GO term. **d**, Heatmaps showing gene selection criteria. Top genes are labelled in red; Asterisk indicates the selected target gene.

To confirm that neuronal activity regulates *Fezf2* expression in L5 ET cells, we reduced the activity of these cells during early postnatal development and performed single-molecule RNA fluorescent in situ hybridization at P10. Kir2.1-expressing ET pyramidal cells exhibited a marked decrease in *Fezf2* mRNA levels compared to controls (Fig. 3a-c). To extend these observations, we transiently increased the excitability of L5 ET neurons using a chemogenetic approach based on Designer Receptors Exclusively Activated by Designer Drugs (DREADDs). In brief, we injected *rAAV_retro_-Cre* in the pons and AAVs encoding excitatory DREADDs (hM3Dq) in the S1 of P1 mice. A single injection of clozapine-*N*-oxide (CNO) at P8 induced the expression of c-Fos, specifically in hM3Dq-expressing neurons (Extended Data Fig. 4). We then injected CNO twice daily between P8 and P10 and examined the expression of *Fezf2* at this stage (Fig. 3d). We found that the expression of *Fezf2* was increased in hM3Dq-infected L5 ET cells in mice treated with CNO compared to vehicle (Fig. 3e, f). These experiments confirmed a cell-autonomous link between the activity of L5 ET pyramidal cells and *Fezf2* expression.

**Fig. 3:**
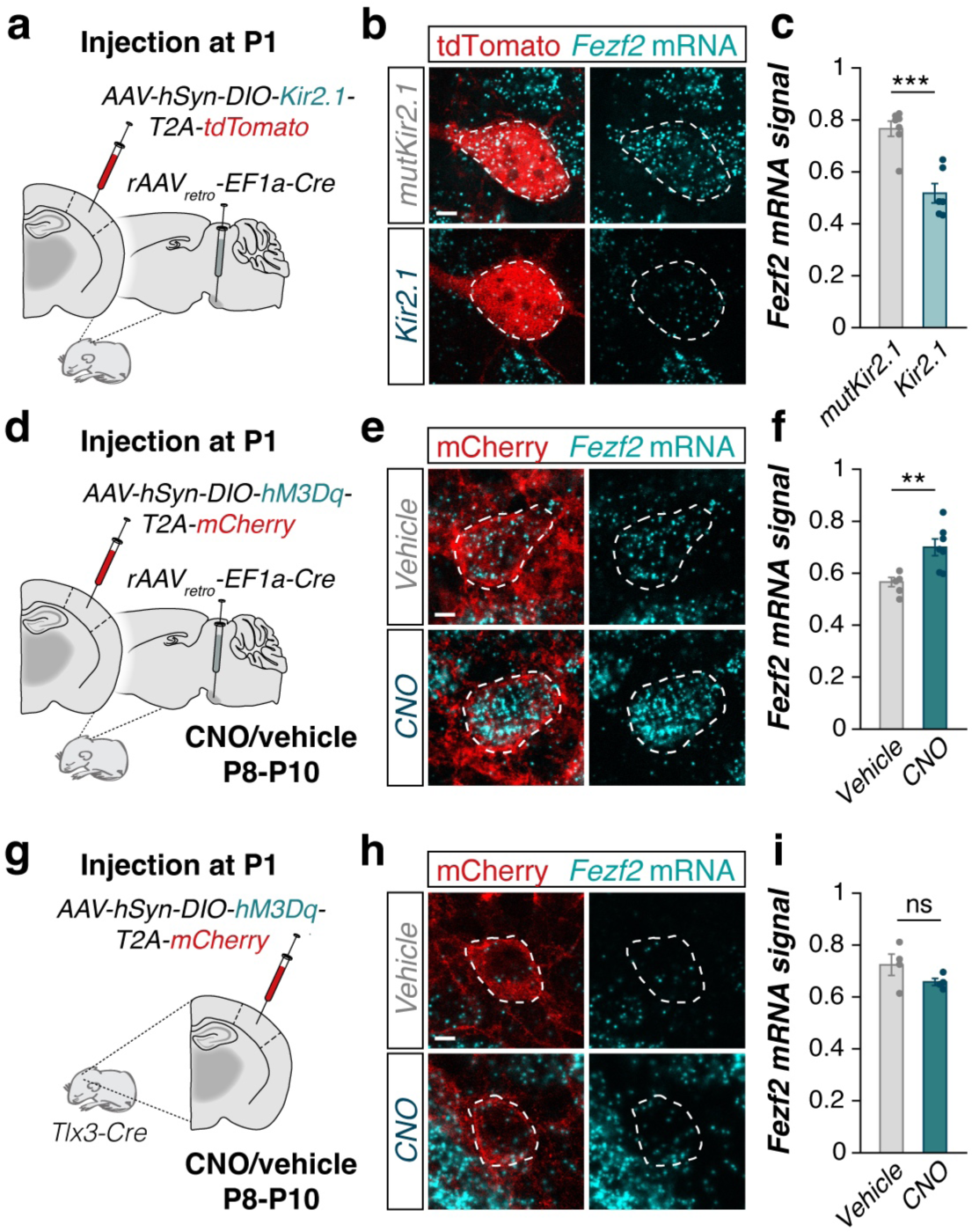
Activity regulation of *Fezf2* is cell-type specific. **a**, Experimental design for decreasing activity with Kir2.1 in ET pyramidal cells. **b**, Confocal images showing *Fezf2* mRNA expression (cyan) in Kir2.1- and mutKir2.1-expressing ET pyramidal cells (tdTomato, red). **c**, Quantification of *Fezf2* mRNA signal in Kir2.1-expressing ET cells compared to controls (mutKir2.1, *n* = 6 animals; Kir2.1, *n* = 6 animals, two-tailed Student’s *t*-test, \*\*\**P* < 0.001). **d**, Experimental design for increasing activity with DREADDs in ET pyramidal cells. **e**, Confocal images showing *Fezf2* mRNA expression (cyan) in hM3Dq-expressing ET pyramidal cells (mCherry, red). **f**, Quantification of *Fezf2* mRNA signal in hM3Dq-expressing ET cells from mice injected with CNO compared to controls (vehicle, *n* = 6 mice; CNO, *n* = 7 animals, two-tailed Student’s *t*-test, \*\**P* < 0.01). **g**, Experimental design for increasing activity with DREADDs in IT pyramidal cells. **h**, Confocal images showing *Fezf2* mRNA expression (cyan) in hM3Dq-expressing IT pyramidal cells (mCherry, red). **i**, Quantification of *Fezf2* mRNA signal in hM3Dq-expressing IT cells from mice injected with CNO compared to controls (vehicle, *n* = 4 mice; CNO, *n* = 4 mice, two-tailed Student’s *t*-test, ns). The dotted line indicates the outline of the soma, drawn from the tdTomato signal. Data are mean ± SEM, ns, not significant. Scale bars, 5 μm (**b**, **e**, **h**).

To investigate whether the activity-dependent expression of *Fezf2* was cell type-specific or could be triggered by changes in activity in other pyramidal cells, we expressed hM3Dq in L5 IT neurons. To this end, we used *Tlx3-Cre* mice, in which Cre expression is specific to L5 IT pyramidal cells^40^. We found that increasing the activity of IT neurons between P8 and P10 does not change the expression of *Fezf2* in these cells (Fig. 3g-3i). This observation suggests that the activity regulation of *Fezf2* is cell type-specific.

### *Fezf2* regulates the assembly and maintenance of inhibition onto ET neurons

FEZF2 plays a critical role in the specification of L5 ET neurons during embryonic development ^35–39^. We found that neuronal activity regulates the expression of *Fezf2* in L5 ET cells during postnatal development, suggesting that this transcription factor may play additional roles in defining important features of these cells long after they have been specified. We hypothesized that FEZF2 might regulate the density of perisomatic PV+ inhibitory inputs received by L5 ET cells in an activity-dependent manner. To test this idea, we engineered Cre-dependent AAVs containing the coding sequence for green fluorescent protein (GFP) and a short hairpin RNA (shRNA) sequence against *Fezf2*. As a control, we used a shRNA sequence against beta-galactosidase (*LacZ*), which does not target mammalian genes^11^. We verified the efficiency of *shFezf2* to downregulate the expression of *Fezf2* in vitro and in vivo (Extended Data Fig. 5a-d). We then injected *rAAV_retro_-Cre* in the pons and *shRNA* AAVs in the S1 at P1 and prepared acute slices from these mice at P15, to allow for sufficient downregulation (Fig. 4a). We found that the downregulation of *Fezf2* leads to a reduction in the frequency of mIPSCs (Fig. 4b, c) and the density of somatic PV+ basket cell boutons contacting L5 ET cells (Fig. 4d, e). These experiments demonstrate that FEZF2 is cell-autonomously required for L5 ET neurons to receive a normal complement of inhibitory synapses from PV+ basket cells.

**Fig. 4:**
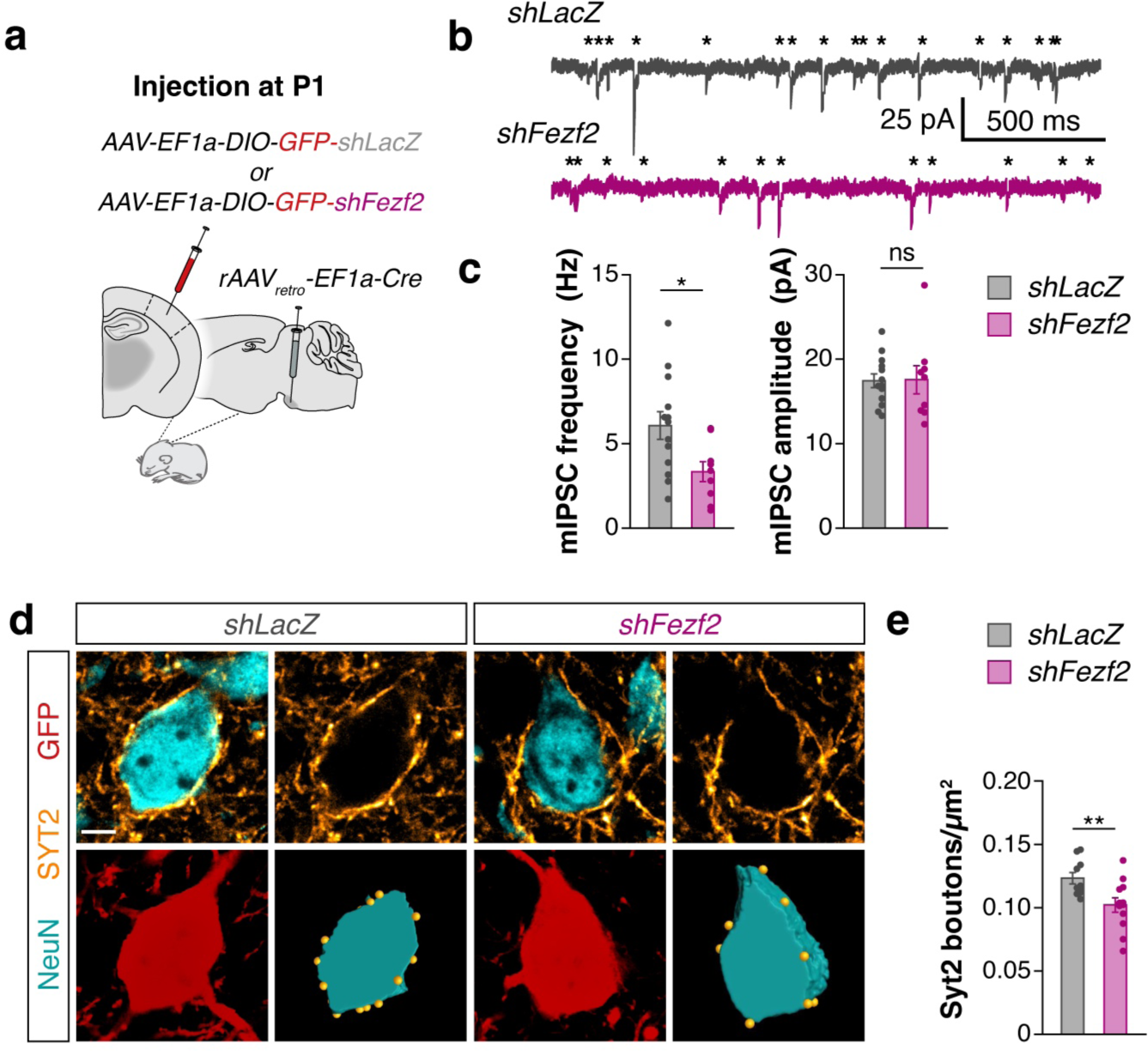
*Fezf2* regulates perisomatic inhibition onto ET pyramidal neurons. **a**, Experimental design of *Fezf2* downregulation. **b**, Example traces showing mIPSCs recorded in ET pyramidal cells infected with either *shLacZ*-*GFP* (top) or *shFezf2*-*GFP* (bottom). Asterisks indicate the onset of mIPSC events. **c**, Quantification of mIPSC frequency (*shLacZ*, *n* = 13 cells from 4 mice; *shFezf2*, *n* = 9 cells from 4 mice, two-tailed Student’s *t*-test, \**P* < 0.05) and amplitude (uninfected, *n* = 13 cells from 4 mice; *shLacZ*, *n* = 13 cells from 4 animals; *shFezf2*, *n* = 9 cells from 4 mice, two-tailed Student’s *t*-test, ns). **d**, Confocal images showing presynaptic Syt2 puncta (yellow), NeuN (cyan) and GFP (red). Bottom right: 3-dimensional Imaris reconstruction of the soma and Syt2 puncta. **e**, Quantification of the density of Syt2 puncta onto the soma (*shLac* Z, *n* = 10 animals; *shFezf2*, *n* = 10 animals, two-tailed Student’s *t*-test, \**P* < 0.05). Data are mean ± SEM, ns, not significant. Scale bar, 5 μm (**d**).

Since *Fezf2* is essential for the specification of L5 ET neurons early in development, postnatal downregulation of *Fezf2* could potentially alter their cell identity, partially reprogramming these cells and thereby secondarily reducing their perisomatic inhibition. To discard this possibility, we investigated whether postnatal downregulation of *Fezf2* affected other key features of L5 ET pyramidal cells. We found no differences in the intrinsic properties, soma size, or axonal projections between *shFezf2*-infected neurons and control cells (Fig. 5a-d). These observations suggested that loss of *Fezf2* does not change the identity of L5 ET pyramidal cells.

**Fig. 5:**
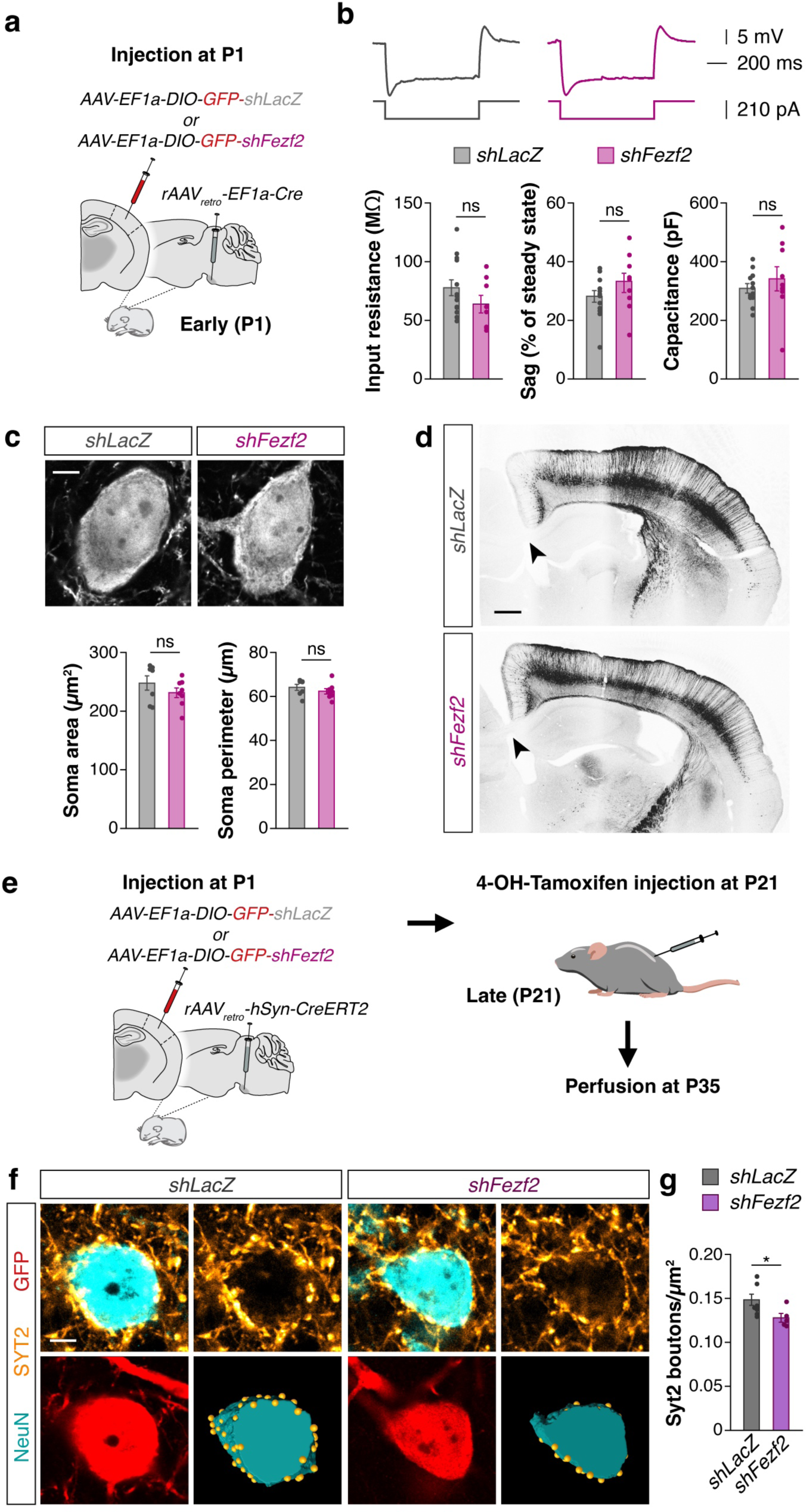
*Fezf2* controls perisomatic inhibition independent of cell type-specific features. **a**, Experimental design of *Fezf2* downregulation. **b**, Example traces showing membrane voltage deflections in response to hyperpolarizing current steps. Bottom: quantification of cell type-characteristic intrinsic properties in *shFezf2* compared to controls: input resistance (*shLacZ*, *n* = 13 from 4 mice; *shFezf2*, *n* = 8 from 4 mice, two-tailed Student’s *t*-test, ns), sag current relative to steady-state (*shLacZ*, *n* = 13 from 4 mice; *shFezf2*, *n* = 9 from 4 mice, two-tailed Student’s *t*-test, ns), and capacitance (*shLacZ*, *n* = 13 from 4 mice; *shFezf2*, *n* = 9 from 4 mice, two-tailed Student’s *t*-test, ns). **c**, Top: Single-plane confocal images of GFP-filled somata of *shLacZ* (left) and *shFezf2*-*infected* (right) cells. Bottom: Quantification of soma size (*shLacZ*, *n* = 7 mice; *shFezf2*, *n* = 8 mice, two-tailed Student’s *t*-test, ns) and perimeter (*shLacZ*, *n* = 7 mice; *shFezf2*, *n* = 8 mice, two-tailed Student’s *t*-test, ns). **d**, Tiled confocal images showing no discernible axonal projections to corpus callosum in *shFezf2.* **e**, Experimental design of late *Fezf2* downregulation. **f**, Single-plane confocal images showing presynaptic Syt2 puncta (yellow), NeuN (cyan) and tdTomato (red). Bottom right: 3-dimensional Imaris reconstruction of the soma and Syt2 puncta. **g**, Quantification of the density of Syt2 puncta onto the soma (*shLacZ*, *n* = 8 mice; *shFezf2*, *n* = 5 animals, two-tailed Student’s *t*-test, \**P* < 0.05). Data are mean ± SEM, ns, not significant. Scale bars, 5 μm (**c**, **f**), 500 μm (**d**).

*Fezf2* continues to be expressed in L5 ET neurons in adulthood^41^, suggesting it may also regulate the maintenance of perisomatic inhibition in these cells at later stages. To explore this possibility, we downregulated *Fezf2* expression at the end of the period of PV+ synapse formation^12^. In brief, we first injected a retrograde AAV encoding an inducible Cre (CreERT2) in the pontine nuclei and either *shFezf2* or *shLacZ* Cre-dependent AAVs in the S1 of wild type mice at P1. We then induced Cre-dependent recombination by the intraperitoneal administration of 4-hydroxytamoxifen (4OH-T) at P21 (Fig. 5e). We confirmed this approach efficiently downregulates *Fezf2* expression by P35 (Extended Data Fig. 5e). We found that, as in the early knockdown experiments, the late downregulation of *Fezf2* knockdown in L5 ET neurons reduces the density of Syt2+ boutons contacting the soma of these cells (Fig. 4f, g). Altogether, our results demonstrated that *Fezf2* is an activity-dependent and cell type-specific transcription factor required for the formation and maintenance of PV+ perisomatic synapses onto L5 ET pyramidal cells.

### Fezf2 induces Cdh22 expression to regulate perisomatic inhibition

Cell-surface molecules are involved in the formation and maintenance of synapses in the central nervous system^42–45^. Notably, the three top-ranked genes downregulated by a reduction in the activity of L5 ET neurons, *Cdh22*, *Cntn6*, and *Igsf21*, encode cell-surface molecules (Fig. 2d). We hypothesized that one of these proteins could function downstream of FEZF2 to regulate PV+ perisomatic inhibition in L5 ET cells. To test this hypothesis, we first assessed whether the expression of *Cdh22*, *Cntn6*, and *Igsf21* is indeed regulated by neuronal activity. Using single-molecule RNA fluorescent in situ hybridization, we observed that the expression of *Cdh22* and *Cntn6* was reduced in Kir2.1-expressing L5 ET cells (Fig. 6a-c). In contrast, *Igsf21* expression was not changed in these experiments (Fig. 6a-6c), suggesting that this gene might be a false-positive in our screen or that activity regulates its translation rather than its transcription. To test whether changes in activity could reversibly affect the expression of *Cntn6* and *Cdh22*, we next increased the activity of ET pyramidal neurons using chemogenetics (Fig. 6d). We observed that increasing neuronal activity in L5 ET cells increased the expression of *Cdh22* but not *Cntn6* (Fig. 6e, f).

**Fig. 6:**
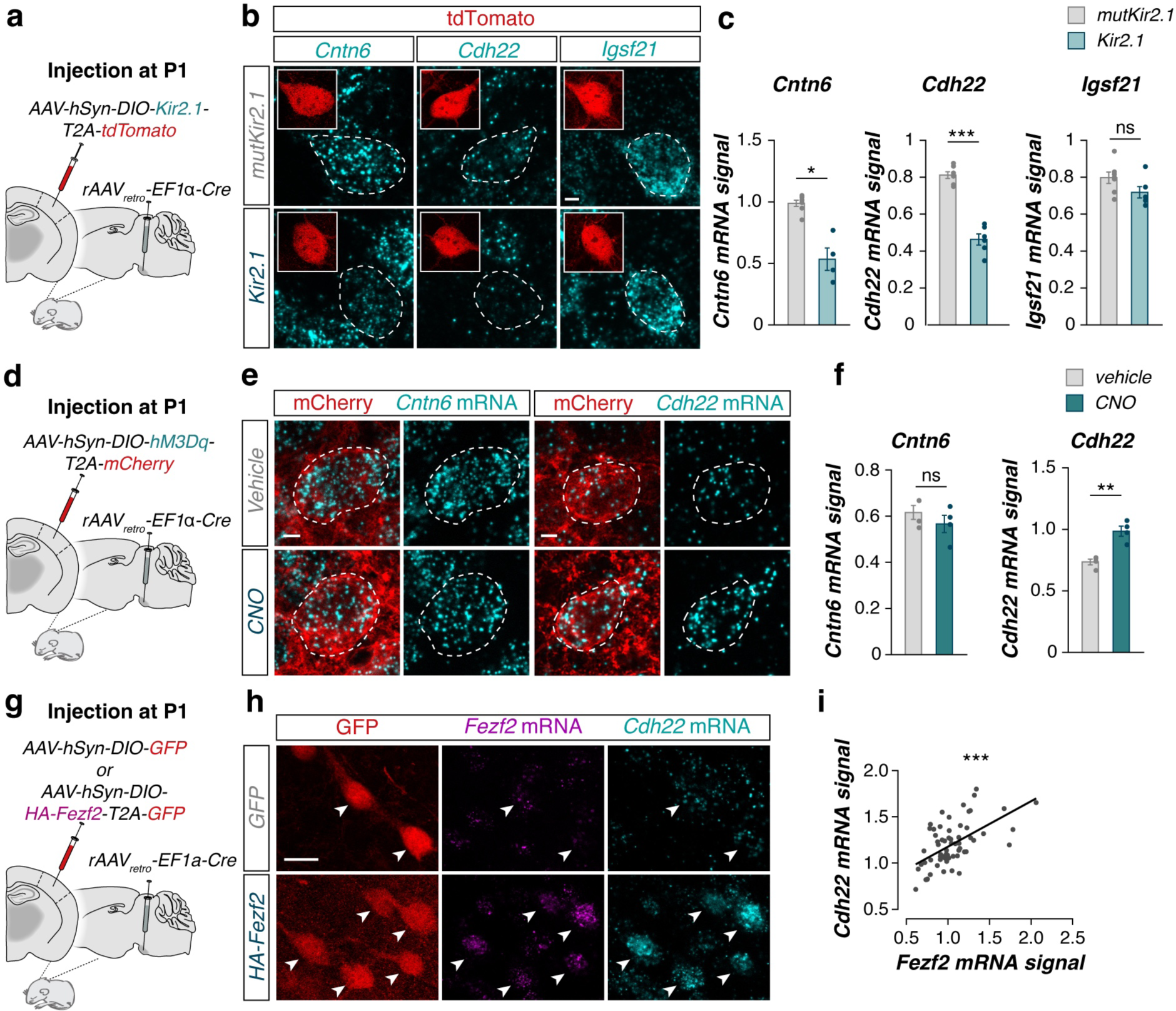
Activity and *Fezf2* regulate *Cdh22* expression. **a**, Experimental design for decreasing activity with Kir2.1 in ET pyramidal cells. **b**, Confocal images showing *Cntn6*, *Cdh22*, and *Igsf21* mRNA expression (cyan) in Kir2.1- and mutKir2.1-expressing ET pyramidal cells (tdTomato, red). **c**, Quantification of *Cntn6*, *Cdh22*, and *Igsf21* mRNA signal in Kir2.1-expressing ET cells compared to controls (*Cntn6: mutKir2.1*, *n = 7 mice; Kir2.1*, *n = 4 mice two*-*tailed Welch’s t*-*test*, *p < 0.05; Cdh22*: mutKir2.1, *n* = 7 mice; Kir2.1, *n* = 6 mice, two-tailed Student’s *t*-test, \*\*\**P* < 0.001; *Igsf21*: mutKir2.1, *n* = 7 animals; Kir2.1, *n* = 6 animals, two-tailed Student’s *t*-test, ns). **d**, Experimental design for increasing activity with DREADDs in ET pyramidal cells. **e**, Confocal images showing *Cntn6* and *Cdh22* mRNA expression (cyan) in *AAV* and *rAAVretro Cre* infected ET pyramidal cells (mCherry, red). **f**, Quantification of *Cntn6* and *Cdh22* mRNA signal in hM3Dq-expressing ET cells from mice injected with CNO compared to controls (Cntn6: *n* = 4 mice; CNO, *n* = 4 mice, two-tailed Student’s *t*-test, ns; Cdh22: vehicle, *n* = 4 mice; CNO, *n* = 4 mice, two-tailed Student’s *t*-test, \*\**P* < 0.01). **g**, Experimental design of *Fezf2* overexpression in ET pyramidal cells. **h**, Confocal images showing *Fezf2* and *Cdh22* mRNA expression (cyan) in *AAV* and *rAAVretro Cre* infected ET pyramidal cells (mCherry, red). **i**, Quantification of the correlation between *Fezf2* and *Cdh22* mRNA signal in ET neurons overexpressing *Fezf2* (n = 64 cells from 1 mouse, simple linear regression: r^2^ = 0.33, *** p < 0.001). Data are mean ± SEM, ns, not significant. Scale bars, 5 μm (**b**, **e**), 20 μm (**h**).

FEZF2 binds closely to the *Cdh22* transcription starting site^38^, which suggests that CDH22 might be a potential direct effector of FEZF2 in mediating the formation of PV+ synapses on the soma of L5 ET neurons. To determine whether FEZF2 regulates *Cdh22* expression, we expressed a Cre-dependent construct encoding HA-tagged *Fezf2* and *Gfp* in L5 ET cells (Fig. 6g) and analyzed the expression of *Cdh22* using in situ hybridization. We observed that *Fezf2* overexpression induces the expression of *Cdh22* (Fig.6h, 6i). Intriguingly, *Cntn6* also seemed modulated by FEZF2, although to a lesser extent than *Cdh22* (Extended Data Fig. 6a-c). Altogether, our findings suggest that the cell-surface molecule *Cdh22* functions downstream of Fezf2 in L5 ET pyramidal cells.

To investigate whether CDH22 is required for the development of PV+ perisomatic inhibition in L5 ET neurons, we engineered a *shRNA* against *Cdh22* mRNA and validated its efficacy in vitro and in vivo (Extended Data Fig. 5f, g). We then performed electrophysiological recordings in L5 ET neurons following the downregulation of *Cdh22* expression (Fig. 7a). We observed that the frequency of mIPSCs was reduced in *shCdh22*-infected cells compared to control neurons (Fig. 7b, c). Consistent with these observations, we found that the downregulation of *Cdh22* also reduced the density of PV+ boutons contacting the soma of L5 ET neurons (Fig. 7d, e). In contrast, the downregulation of *Cntn6* did not affect the frequency or amplitude of mIPSCs in L5 ET pyramidal cells (Extended Data Fig. 6d-i). These findings revealed that an activity-dependent molecular program involving FEZF2 and Cdh22 regulates the development of inhibitory connectivity between PV+ interneurons and L5 ET pyramidal cells.

**Fig. 7:**
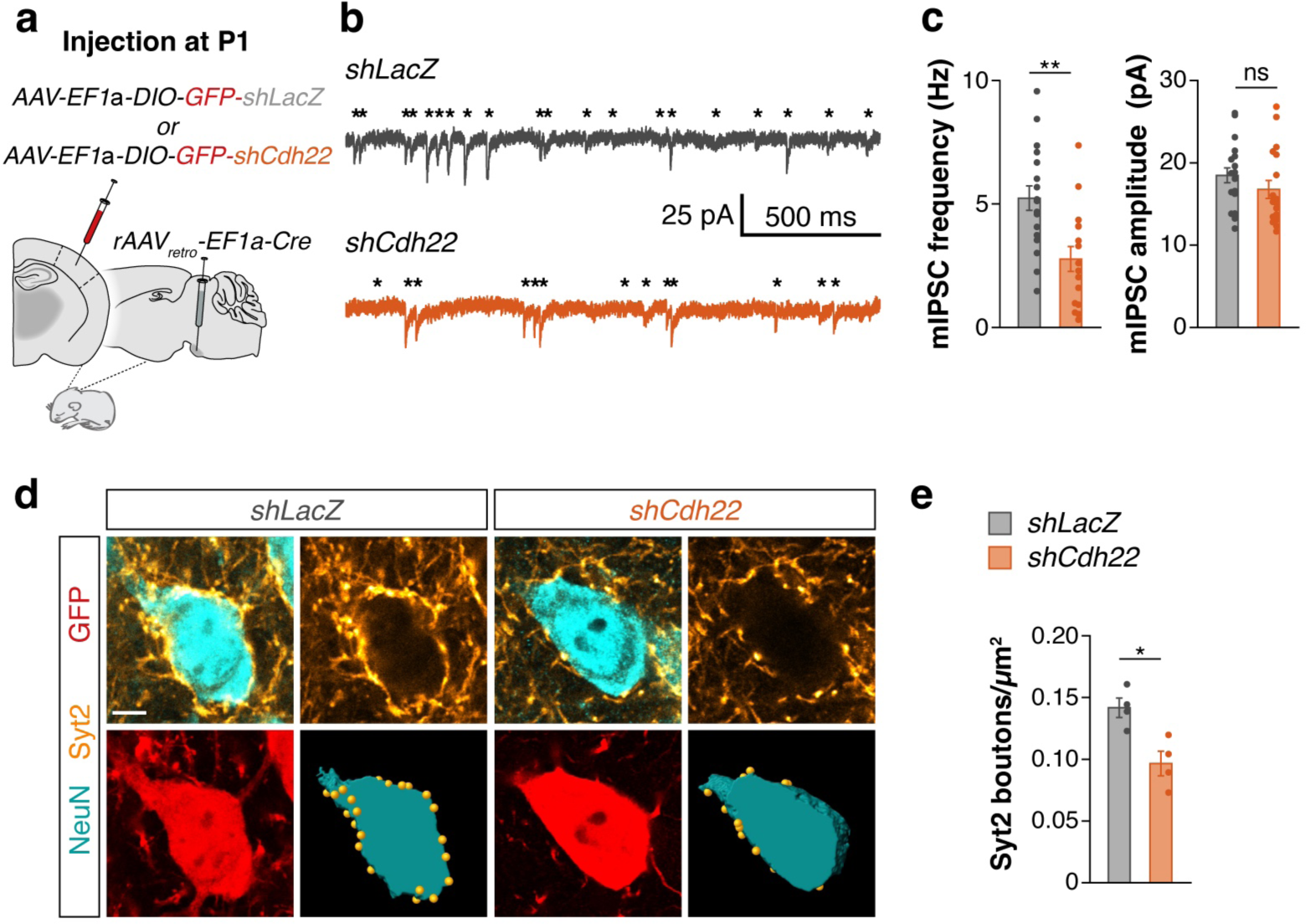
*Cdh22* controls perisomatic inhibition onto ET pyramidal neurons. **a**, Experimental design of *Cdh22* downregulation. **b**, Example traces showing mIPSCs recorded in ET pyramidal cells infected with either *shLacZ* (top) or *shCdh22* (bottom) from P15-17 mice. Asterisks indicate the onset of mIPSC events. **c**, Quantification of mIPSC frequency (*shLacZ*, *n* = 18 cells from 9 mice; *shCdh22*, *n* = 16 cells from 7 mice, two-tailed Student’s *t*-test, \*\**P* < 0.01) and amplitude (shLacZ, *n* = 19 cells from 9 mice; *shCdh22*, *n* = 18 cells from 7 mice, two-tailed Student’s *t*-test, ns). **d**, Confocal images showing presynaptic Syt2 puncta (yellow), NeuN (cyan) and GFP (red). Bottom right: 3-dimensional Imaris reconstruction of the soma and Syt2 puncta. **e**, Quantification of the density of Syt2 puncta close to the soma (*shLacZ*, *n* = 4 mice; *shFezf2*, *n* = 4 mice, two-tailed Student’s *t*-test, \**P* < 0.05). Data are mean ± SEM, ns, not significant. Scale bar, 5 μm (**d**).

## Discussion

In the adult cerebral cortex, neuronal networks exhibit remarkable flexibility to integrate fluctuations in neuronal activity during ongoing environmental changes^17^. Pyramidal cells maintain their firing rates by regulating the level of inhibition they receive, which involves activity-dependent regulation of gene expression (Yap and Greenberg, 2018). In this process, broadly expressed IEGs trigger the expression of effector factors in a cell type-specific manner^22, 23^. Our study reveals that activity-dependent mechanisms are already at play during early postnatal development to regulate the functional integration of pyramidal cells into nascent cortical networks. We demonstrate that *Fezf2*, a selector gene involved in specifying the identity of L5 ET cells in the embryo, is regulated postnatally by neuronal activity exclusively in these neurons. We also found that activity-dependent changes in *Fezf2* expression sculpt the formation of PV+ synapses onto ET pyramidal cells via the cell-surface molecule CDH22. These experiments uncovered a cell type-specific transcriptional mechanism that sculpts the development of cortical circuitries in an activity-dependent manner. Our findings suggest that cell type-specific intermediate transcription factors such as FEZF2 provide cell-type specificity to the activity-dependent regulation of cortical wiring.

Our findings revealed that overexpression of *Fezf2* increases the expression of *Cdh22* and *Cntn6* transcripts, suggesting that both cell-surface proteins are downstream of *Fezf2*. Experiments using chromatin immunoprecipitation followed by deep sequencing of progenitor cells from neurospheres identified direct FEZF2 binding sites close to the transcription start site of *Cdh22* but not *Cntn6*^38^. These observations suggest that *Cntn6* is probably not directly regulated by FEZF2. Our experiments demonstrated that FEZF2 and neuronal activity regulate *Cdh22* expression and that *Cdh22* knockdown decreases the density of PV+ somatic synapses contacting L5 ET pyramidal cells. These findings suggest that *Cdh22* may be a downstream effector of FEZF2 in the regulation of inhibitory synapses. CDH22 can form homophilic and heterophilic interactions with other cadherin family members^46^. Of these, *Cdh7* expression is enriched in PV+ basket cells during postnatal development^11, 47^, suggesting a potential direct mechanism for the involvement of CDH22 in PV+ basket cell synapse formation onto ET neurons. We also observed that the downregulation of *Cntn6* does not affect the density of inhibitory inputs targeting L5 ET pyramidal cells. Thus, *Cntn6* may play a different role in shaping the maturation of L5 ET neurons.

The development of inhibitory synapses in the cerebral cortex spans an extended period of postnatal development^11, 13, 14^. Although the density of PV+ basket cell inputs to pyramidal cells increases 3-fold around P9, somatic synapses continue to grow steadily until P28^14^. We have recently demonstrated that *Cdh13* is required for the cell-specific targeting of L5 ET pyramidal cells by PV+ basket cells^12^. Therefore, CDH13 may be part of a genetically imprinted code for cell-specific target selection in ET pyramidal cells, which receive progressively more excitatory inputs during early postnatal development. This increase in excitatory drive would then trigger the activity-regulated expression of *Fezf2* and its downstream effector *Cdh22* to recruit PV+ basket cell inputs and shape this circuitry towards their maturity.

The balance of excitation and inhibition (E/I balance) in the cerebral cortex is essential for the stabilization of neural circuit function^48^, including learning and memory^49, 50^. To maintain an appropriate E/I balance in response to changes associated with sensory experience, pyramidal cells recruit inhibitory input in an activity-dependent manner^18, 19, 21^. This is mediated by the rapid initial expression of IEGs, most of which encode transcription factors that regulate a second wave of LRGs^22^. While most IEGs encode transcription factors such as FOS, JUN, and NPAS4^22^, cell type-specific LRGs often encode molecules that are secreted or localized to the synapse^51–56^. These molecular programs generate the foundation for structural and functional modifications at the synapses. In the last decade, it has become clear that activity-regulated gene expression is not homogeneous across neuron types. Instead, while a common set of IEGs can be expressed across all neurons, the complement of LRGs that a neuron expresses in response to IEG induction often varies from one neuron type to another^23, 54, 57, 58^. This cell-type specificity is hypothesized to come about through the combined action of commonly expressed activity-dependent transcription factors and cell type-specific transcription factors that are not activity-regulated at the same enhancers^23, 59^, but such factors have not been identified.

We have shown that activity regulates the expression of *Fezf2* in ET cells but not in other pyramidal cells. To our knowledge, this is the first example of a cell type-specific transcription factor regulated by activity. IEGs, such as FOS, are expressed in L5 ET neurons ^60, 61^. As shown for other pyramidal cells^18, 21^, FOS may also adjust the inhibitory wiring in L5 neurons. An appealing hypothesis is that in addition to directly triggering known general effector molecules like BDNF, IGF1, and SCG2^21, 59^, IEGs also regulate intermediate transcription factors that confer specificity to the wiring of specific populations of neurons, such as *Fezf2*. Although the existence of such intermediate transcription factors has been hypothesized, they have not been identified, likely due to a bias towards effector genes in the selection of candidates. Interestingly, the transcription factor Etv1/Er81 is required for the development of specific proprioceptive neuron connectivity in the spinal cord^62^ and is an activity-regulated gene PV interneurons^63^. It remains to be explored whether Etv1 plays a role similar to FEZF2 in other neurons.

*Fezf2* is a selector gene for L5 ET pyramidal cells, as its removal in embryos leads to a loss of cell type identity^35–39^. We found that postnatal *Fezf2* expression controls inhibitory synapse formation in L5 ET pyramidal cells but its loss does not impact cell identity, which contrasts the function of this transcription factor at embryonic stages. Previous studies suggested deep layer cell identity is locked in the first postnatal week^64, 65^. Interestingly, visual experience regulates upper-layer cell identity but not deep-layer cell identity^66^. Although our knockdown strategy might not abolish FEZF2 function completely, the level of downregulation we achieve in our experiments is enough to disrupt the inhibition received by L5 ET pyramidal cells without affecting other prominent features of these neurons. This observation, therefore, indicates that the decrease in inhibition is caused by the downregulation of *Fezf2* in L5 ET neurons. The dual role of *Fezf2* during development is likely the result of a change in the target genes bound by *Fezf2* at different developmental stages. Consistent with this idea, *Fezf2* downregulation in adult mice reveals changes in a markedly different set of genes than during development^38, 41^. Our findings revealed that the expression of *Fezf2* is regulated by neuronal activity postnatally. Other transcription factors seem to be regulated by activity during postnatal stages. For example, *Cpg15* gene expression is only activity-regulated during synapse formation^55, 67–70^. Similarly, *Satb1* is also expressed during early development, but its expression is only regulated in an activity-dependent manner postnatally^51^. *Fezf2* may also have a similar two-stage regulation, activity-independent before birth and activity-dependent after birth.

It is tempting to speculate that cell type-specific transcription factors regulated by IEGs may function as master regulators coordinating multiple transcription programs. Master regulators would differ among neurons to better fine-tune the circuitry depending on cell type-specific demands. In this scenario, what better choice for a neuron than recycling a selector gene, an already cell-type identity and specific transcription factor and utilizing it postnatally in an activity-dependent manner. Future investigations should aim to elucidate whether other neurons use cell type-specific activity-regulated transcription factors or if this is a unique feature of ET pyramidal cells.

## Data availability

Sequencing data have been deposited at the National Center for Biotechnology Information BioProjects Gene Expression Omnibus (GEO) and will be available upon publication. All constructs generated in this study will be available upon publication.

## Author contributions

T.K. and B.R. conceived and designed the study. T.K. performed most of the experiments and analyses. S.S. and H.Y. performed additional experiments and analyses. T.K. designed and produced the molecular tools, with the help of T.G. and P.M.; T.G. and P.M. helped with the *in vitro* experiments and the AAV virus production. S.S., T.G., P.M. helped with the tissue preparation and immunohistochemistry. T.K. and B.R. wrote the manuscript.

## Acknowledgements

We are thankful to L. Doglio, F. Sanchez-Roman, S. Sanalidou and T. Garces for technical assistance and lab support, and to Ian Andrew for mouse management. We thank the CRG Genomics Core Facility of Barcelona for conducting RNA sequencing. Finally, we are grateful to O. Marín, N. Flames, L. Andreae, M. Selten, J. Jézéquel and G. Condomitti for critical reading of the manuscript and to all members of the Rico and Marín laboratories for stimulating discussions and ideas. This project was supported by grants from Wellcome Trust (202758/Z/16/Z) and European Union’s Horizon 2020 research and innovation program (AIMS -2- TRIALS, 777394) to B.R.

---

**Extended Data Fig. 1:**
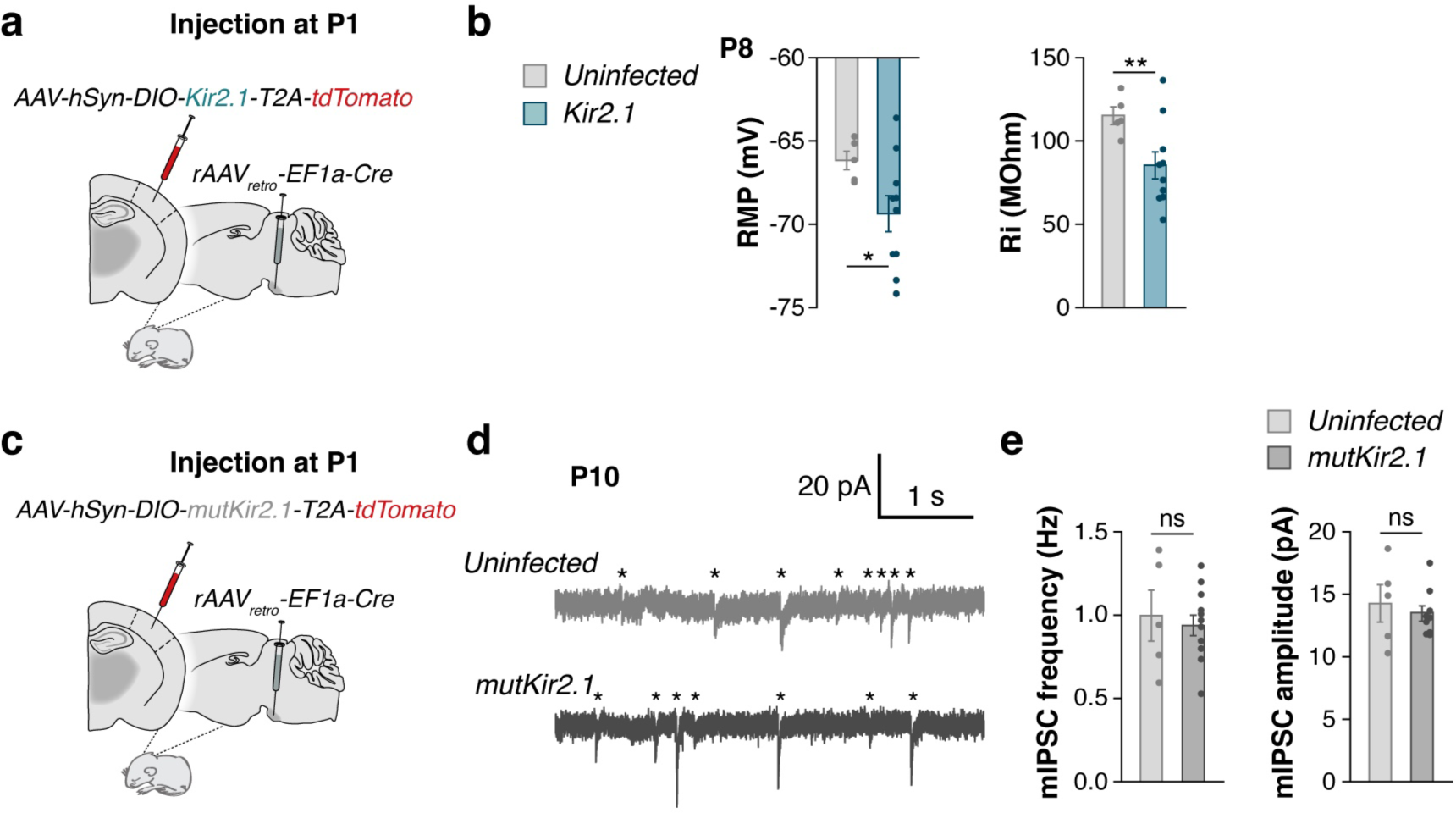
Expression of Kir2.1 decreases excitability of L5 ET pyramidal cells. **a**, Experimental design for decreasing activity with Kir2.1. **b**, Quantification of resting membrane potential (RMP), input resistance (Ri) and rheobase (uninfected, *n* = 5 cells from 2 mice; Kir2.1, *n* = 10 cells from 2 mice, two-tailed Student’s *t*-test, \**P* < 0.05, \*\**P* < 0.01, \*\*\**P* < 0.001). **c**, Experimental design. **d**, Example traces of mIPSCs recorded from mutKir2.1-infected and uninfected cells. Asterisks indicate the onset of mIPSC events. **e**, Quantification of mIPSC frequency (uninfected, *n* = 5 cells from 3 mice; mutKir2.1, *n* = 10 cells from 4 mice, two-tailed Student’s *t*-test, ns) and amplitude (uninfected, *n* = 5 cells from 3 mice; mutKir2.1, *n* = 9 cells from 3 mice, two-tailed Student’s *t*-test, ns). Data are mean ± SEM, ns, not significant.

**Extended Data Fig. 2:**
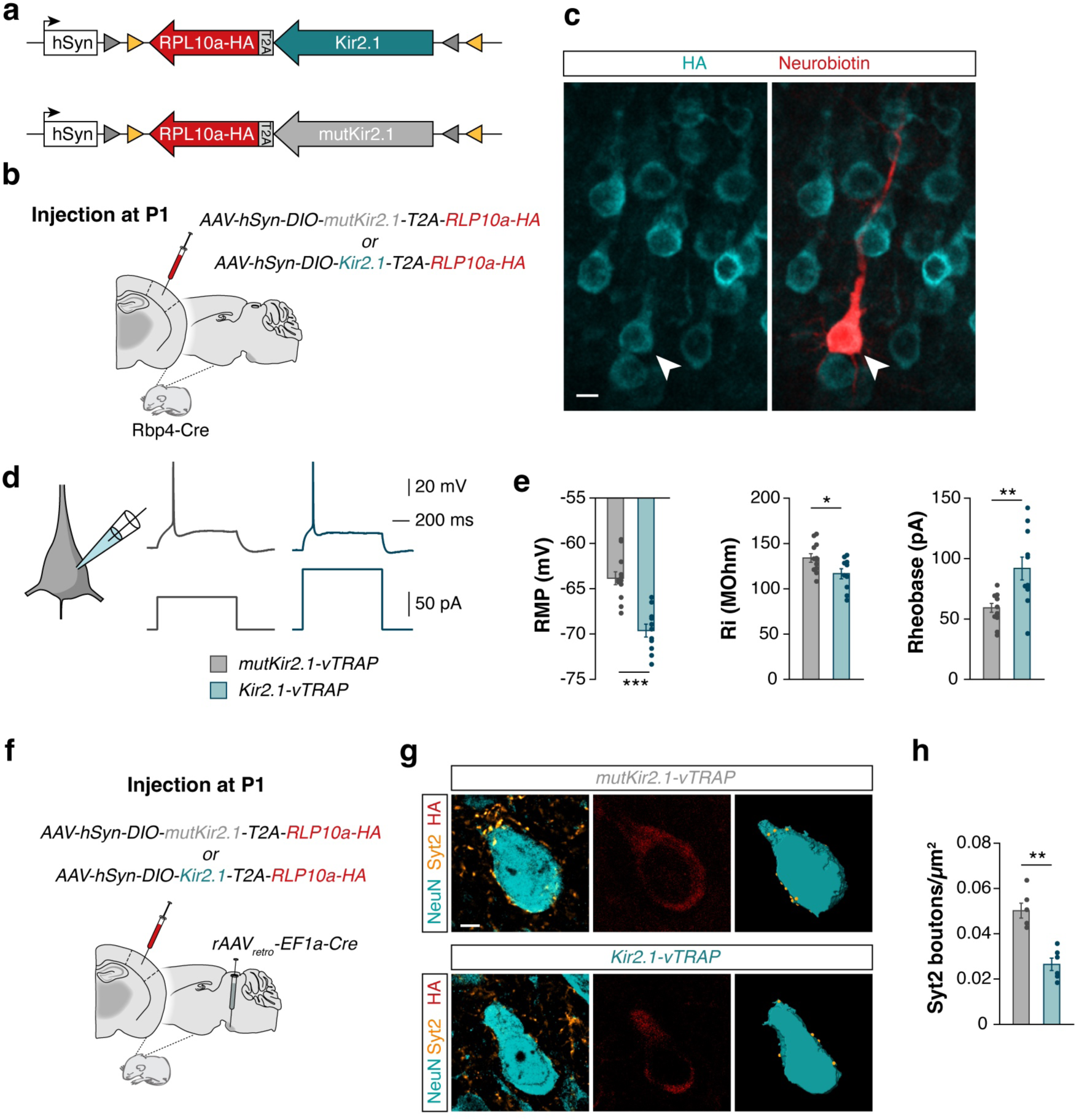
Kir2.1-vTRAP is functional and decreases the density of perisomatic SYT2+ boutons. **a**, Schematic representation of the Cre-dependent Kir2.1-T2A-Rpl10a-HA (Kir-vTRAP) construct. **b**, Experimental design. **c**, Apotome image showing immunohistochemical staining against HA and fluophore-conjugated streptavidin staining of neurobiotin in a recorded cell (arrowhead). **d**, Example traces of the first current injection to elicit an action potential in mutKir2.1-vTRAP and Kir2.1-vTRAP-infected cells. **e**, Quantification of resting membrane potential (RMP; mutKir2.1-vTRAP, *n* = 12 cells from 2 mice; Kir2.1-vTRAP, *n* = 11 cells from 2 mice, two-tailed Student’s *t*-test, \*\*\**P* < 0.001), input resistance (Ri; mutKir2.1-vTRAP, *n* = 12 cells from 2 mice; Kir2.1-vTRAP, *n* = 10 cells from 2 mice, two-tailed Student’s *t*-test, \**P* < 0.05) and rheobase (mutKir2.1-vTRAP, *n* = 12 cells from 2 mice; Kir2.1-vTRAP, *n* = 11 cells from 2 mice, two-tailed Student’s *t*-test, \*\**P* < 0.01). **f**, Experimental design. **g**, Single-plane confocal images showing presynaptic Syt2 puncta (yellow), NeuN (cyan) and HA (red). Right panel: 3-dimensional Imaris reconstruction of the soma and Syt2 puncta. **h**, Quantification of the density of Syt2 puncta close to the somata of cell infected with mutKir2.1-vTRAP or Kir2.1-vTRAP (mutKir2.1-vTRAP, *n* = 6 mice; Kir2.1-vTRAP, *n* = 6 mice, two-tailed Student’s *t*-test, \*\**P* < 0.01). Data are mean ± SEM, ns, not significant. Scale bars, 100 μm (**b**), 5 μm (**f**).

**Extended Data Fig. 3:**
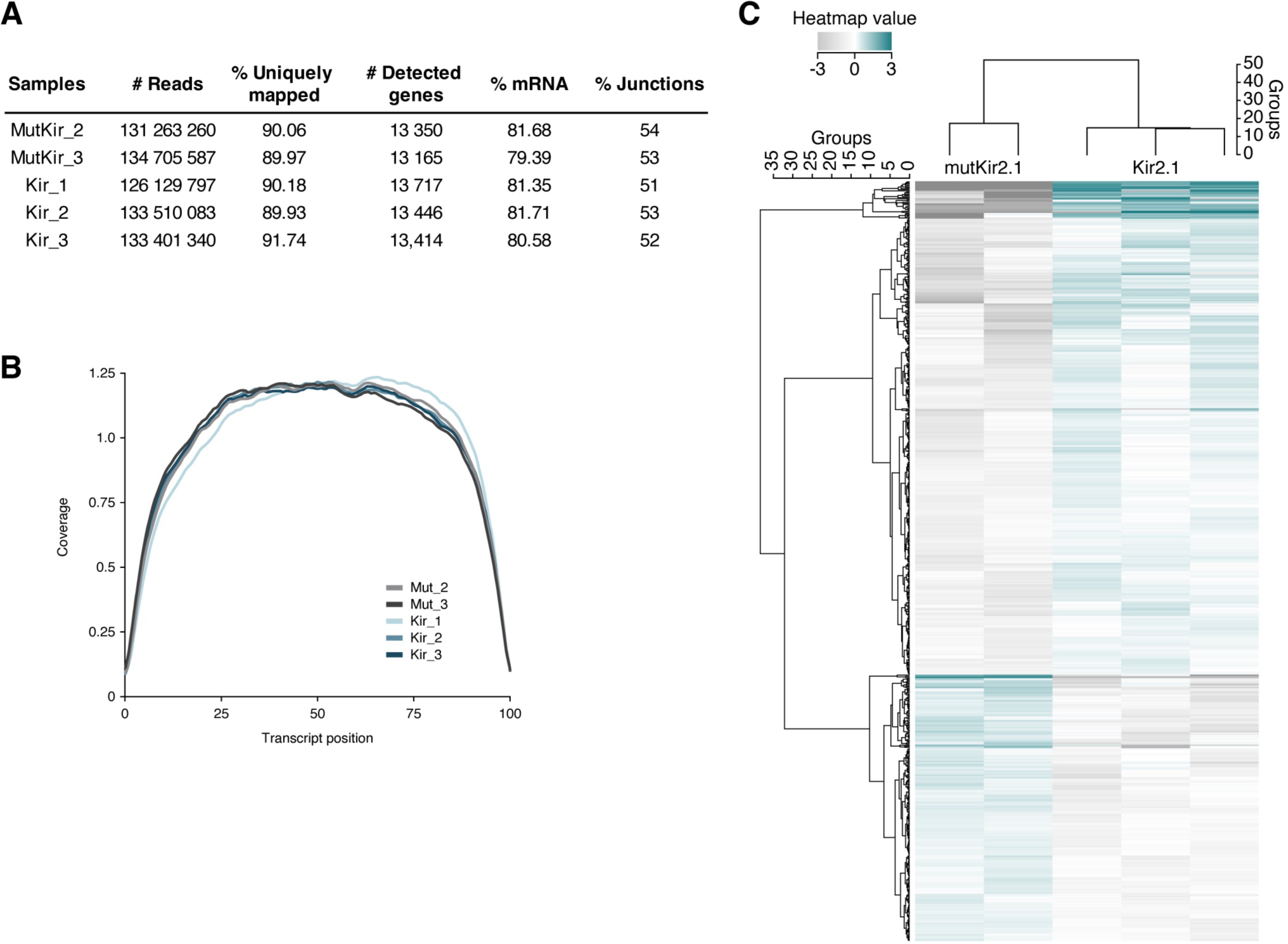
RNA-sequencing quality control and analysis. **a**, FastQC analysis of RNA sequencing data for each replicate, showing the total number of reads, the percentage of reads uniquely mapped to the reference genome, the number of detected genes, the percentage of mRNA and the percentage of reads mapped to exon-exon junctions. **b**, 5’ to 3’ coverage plot showing the percentage of read bases across the transcript length. **c**, Heatmaps showing Deseq2 normalized counts from each replicate of differentially expressed genes.

**Extended Data Fig. 4:**
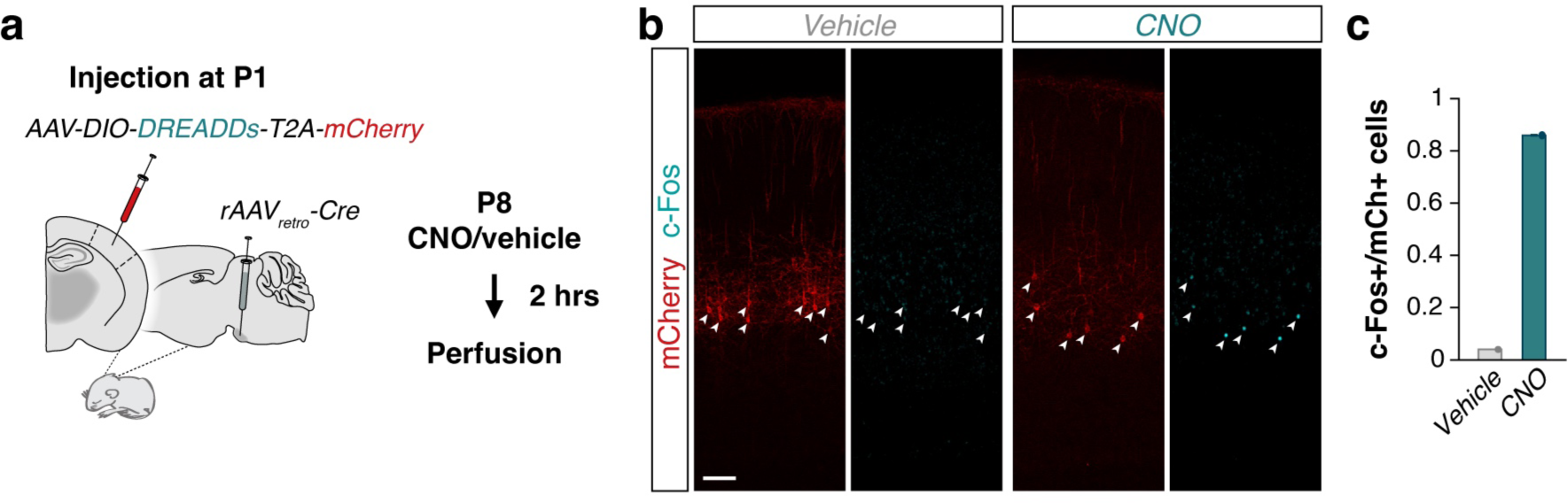
Validation of hM3Dq function. **a**, Experimental setup. Animals were perfused two hours after a single injection of CNO or vehicle. **b**, Single-plane confocal images showing hM3Dq-expressing cells (arrowheads; mCherry, red) and CFOS (cyan). **c**, Quantification of the fraction of hM3D1-expressing cells expressing CFOS after injection of CNO (86%, 2 animals) or vehicle (4%, 1 animal).

**Extended Data Fig. 5:**
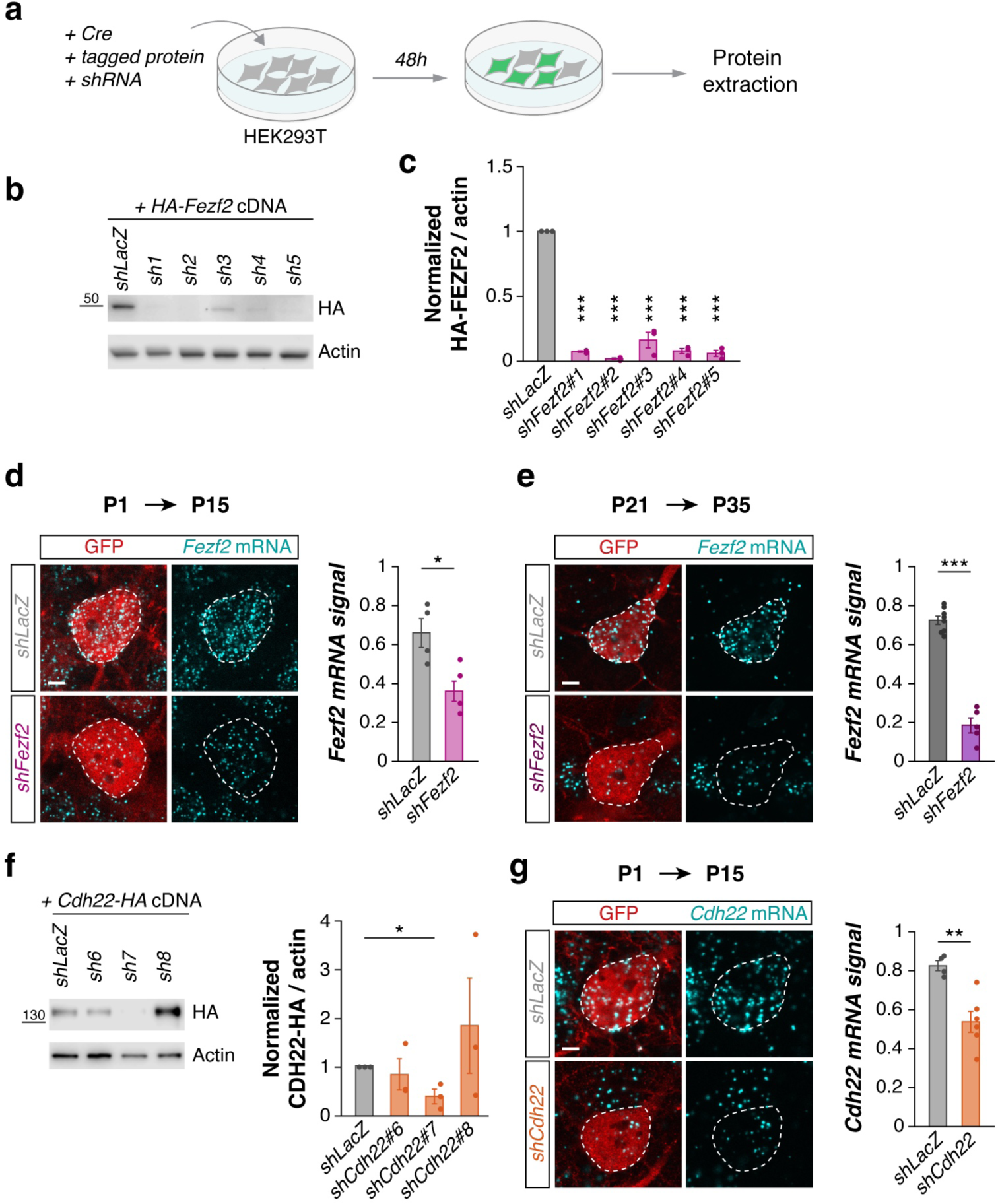
Testing and validation of *Fezf2* and *Cdh22 shRNA* sequences. **a**, Experimental setup for *in vitro* testing of *shRNA* sequences. **b**, Western blot of HA-tagged FEZF2 purified from HEK cells transfected with *HA*-*Fezf2* cDNA and plasmids encoding either *shLacZ* or one of five *shRNAs* against *Fezf2*. **c**, Quantification of protein signal normalized to actin, relative to controls transfected with *shLacZ* (n = 3 wells per sequence, Bonferroni-corrected two-tailed Student’s *t*-test, \*\*\**P* < 0.001). **d**, *Left:* confocal images showing *Fezf2* RNAscope signal (cyan) and tdTomato (red) signal in the *Fezf2* knockdown. *Right*: quantification of *Fezf2* RNAscope signal in the *Fezf2* knockdown compared to controls (*shLacZ*, *n* = 4 mice; *shFezf2*, *n* = 5 mice, two-tailed Student’s *t*-test, \**P* < 0.05). **e**, *Left*: Confocal images showing *Fezf2* RNAscope signal (cyan) and tdTomato (red) signal in the *Fezf2* knockdown from P21 to P35. The dotted line indicates the outline of the soma, drawn from tdTomato signal. Scale bar 5 µm. *Right*: Quantification of *Fezf2* RNAscope signal in the *Fezf2* knockdown from P21 to P35 compared to controls (*shLacZ*, *n* = 8 mice; *shFezf2*, *n* = 5 mice, two-tailed Student’s *t*-test, \*\*\**P* < 0.001). **f**, *Left*: Western blot of HA-tagged FEZF2 purified from HEK cells transfected with *Cdh22*-*HA* cDNA and plasmids encoding either *shLacZ* or one of five *shRNAs* against *Cdh22*. *Right*: quantification of protein signal normalized to actin, relative to controls transfected with *shLacZ* (n = 3 wells per sequence, Bonferroni-corrected two-tailed Student’s *t*-test, \**P* < 0.05). **g**, *Left*: Confocal images showing *Cdh22* RNAscope signal (cyan) and tdTomato (red) signal in the *Cdh22* knockdown. The dotted line indicates the outline of the soma, drawn from tdTomato signal. Scale bar 5 µm. *Right*: quantification of *Cdh22* RNAscope signal in the *Cdh22* knockdown compared to controls (*shLacZ*, *n* = 4 mice; *shCdh22*, *n* = 6 mice, two-tailed Welch’s *t*-test, \*\**P* < 0.01). The dotted line indicates the outline of the soma, drawn from tdTomato signal. Data are mean ± SEM, ns, not significant. Scale bars, 5 μm (**d**, **e**, **g**).

**Extended Data Fig. 6:**
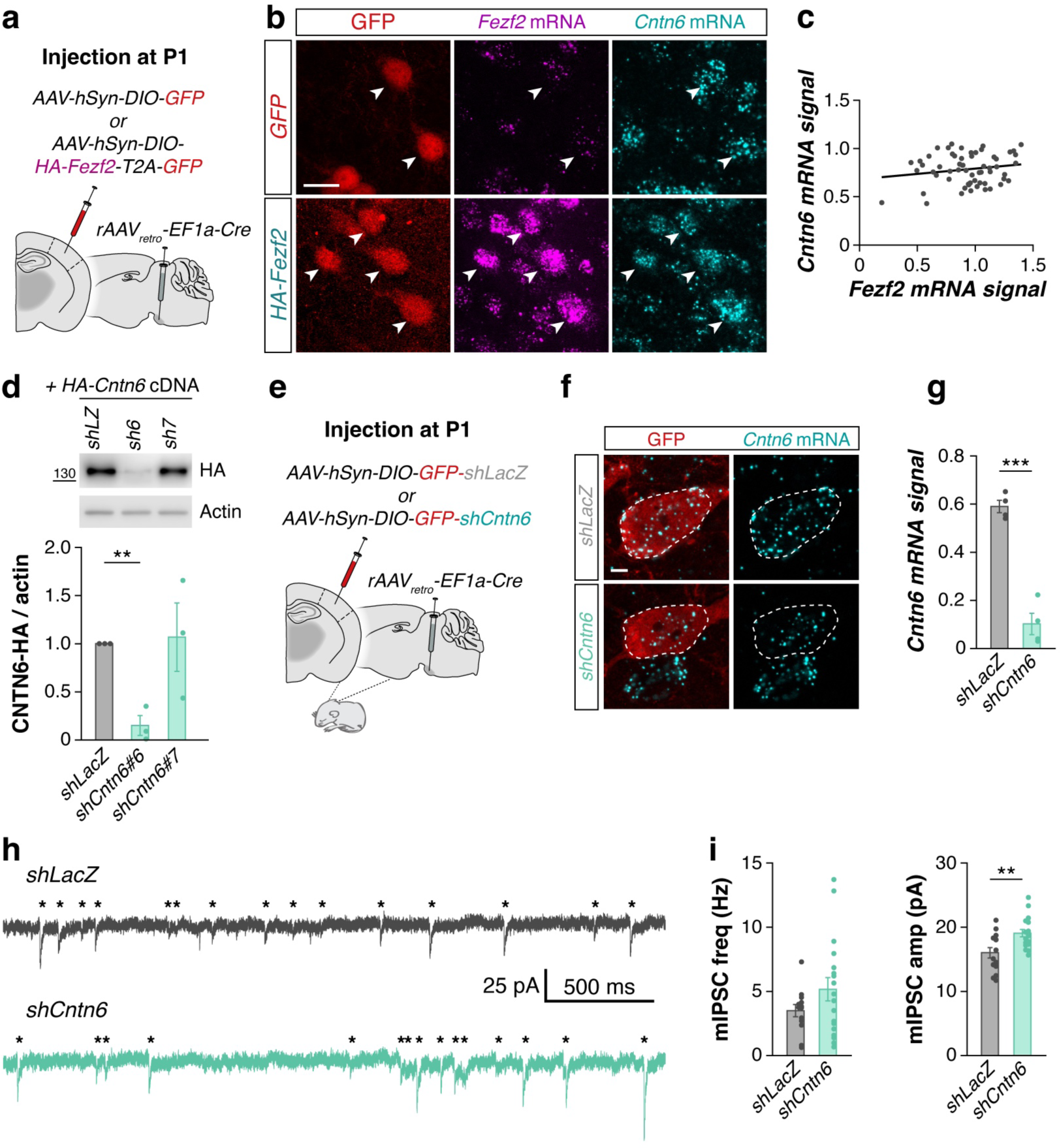
*Cntn6* does not regulate inhibition downstream of *Fezf2*. **a**, Experimental design of *Fezf2* downregulation. **b**, Confocal images showing GFP reporter (red), *Fezf2* mRNA (magenta) and *Cntn6* mRNA (cyan signal) in *Fezf2*-overexpressing and GFP-expressing control cells. **c**, Correlation between *Fezf2* and *Cntn6* mRNA signal in ET neurons overexpressing *Fezf2* (n = 60 cells from 1 mouse, simple linear regression: r^2^ = 0.03, ns). **d**, *In vitro* testing of shRNA sequences. *Left*: Western blot of HA-tagged Cntn6 purified from HEK cells transfected with Cntn6-HA cDNA and plasmids encoding *shLacZ* or one of two *shRNAs* against Cntn6. *Right*: quantification of protein signal normalized to actin, relative to controls transfected with *shLacZ* (n = 3 wells per sequence, Bonferroni-corrected two-tailed Student’s *t*-test, \*\**P* < 0.01). **e**, Experimental design of *Cntn6* overexpression in ET pyramidal cells. **f**, *In vivo* validation of downregulation of endogenous *Cntn6* expression by single-molecule FISH. *Left*: confocal images showing *Cntn6* mRNA (cyan) and GFP (red). **g**, *Right*: quantification of the downregulation of *Cntn6* between *shCntn6* and controls (*shLacZ*, *n* = 4 mice; *shCntn6*, *n* = 4 mice. \*\*\**P* < 0.001). **h**, Example traces showing mIPSCs recorded in ET-PNs infected with either *shLacZ* (top) or *shCntn6* (bottom) from P15-17 mice. Asterisks indicate the onset of mIPSC events. **i**, Quantification of mIPSC frequency (*shLacZ*, *n* = 13 cells from 9 mice; *shCntn6*, *n* = 18 cells from 9 mice, two-tailed Student’s *t*-test, p = 0.16) and amplitude (*shLacZ*, *n* = 14 cells from 9 mice; *shCntn6*, *n* = 18 cells from 9 mice, two-tailed Student’s *t*-test, \*\**P* < 0.01). The dotted line indicates the outline of the soma drawn from the tdTomato signal. Data are mean ± SEM, ns, not significant. Scale bars, 20 μm (**b**), 5 μm (**f**).

## Methods

### Animals

All animal procedures were approved by the ethical committee (King’s College London) and conducted in accordance with European regulations and under licence from the UK Home Office following the Animals (Scientific Procedures) Act 1986. Similar numbers of male and female mice were used in all experiments. Controls were always littermates of experimental animals. Post-weaning, animals were housed in groups of up to five littermates and maintained under standard, temperature-controlled laboratory conditions. Mice were kept on a 12:12 light/dark cycle and received water and food *ad libitum*. Wild-type animals were C57BL/6J (Charles River Laboratories). *Rbp4^Cre^ and Tlx3*^*Cre* 40^ were maintained in a C57BL/6J background.

### CNO treatment

Clozapine N-oxide (CNO; Tocris #4936) was dissolved in 0.5% Dimethyl Sulfoxide (DMSO, Sigma #D0418) in 0.9% saline and stored at -20 °C. Vehicle control was 0.5% DMSO in 0.9% saline was used. Mice were treated with CNO for 48 h starting at P8: two injections per day (morning, between 8 and 9:30 a.m. and evening, between 6 and 8 p.m.) for a total of 5 injections. Animals were then perfused starting 2 hours after the final injection on P10.

To verify the function of hM3Dq and CNO, animals received a single injection of CNO or vehicle and were perfused 2 hours later. Immunohistochemistry was performed to assess the expression of CFOS.

### 4-hydroxytamoxifen treatment

Animals expressing inducible Cre (CreERT2) received a single injection of 4-hydroxytamoxifen (4-OHT; Sigma #H6278) on P21. On the day of injection, 4-OHT was dissolved in ethanol at 55 °C at 100 mg/ml, and further diluted in sunflower seed oil (Sigma #S5007) to a final concentration of 10 mg/ml. 4-OHT was administered via intraperitoneal injection at 50mg/kg.

### Plasmid design

The coding sequences for Kir2.1-T2A-tdTomato and mutKir2.1-T2A-tdTomato (gifts from Massimo Scanziani, Addgene #60661 and #60644, respectively) were subcloned into an hSyn-DIO backbone. *Kir2.1-T2A-Rpl10a-HA* and *mutKir2.1-T2A-Rpl10a-HA* inserts were constructed by GeneArt (Invitrogen) and subcloned into an hSyn-DIO backbone.

The coding sequences of *Fezf2* (NM_080433.3), *Cdh22* (NM_174988.3) and *Cntn6* (NM_017383.3) containing an HA tag at either the N-terminus (*Fezf2* and *Cntn6*) or C-terminus (*Cdh22*), followed by a T2A peptide sequence and GFP, were produced and subcloned into a pcDNA 3.1(+) vector by GeneArt (Invitrogen). *Fezf2-HA-T2A-GFP* was subsequently subcloned into a hSyn-DIO backbone for viral production.

### Short-hairpin RNA design

Using *pAAV-EF1a-DIO-mCherry* described before, the coding sequence for mCherry was replaced with the coding sequence for GFP. The ssDNA primers to generate the shRNAs were either obtained using the Block-it RNAi web tool (Thermo Scientific) or designed by hand and were as follows:

*shFezf2#1* (fwd: CTA GGC GGC TCT CTC TAC TAC TTC ACC TGA CCC ATG AAG TAG TAG AGA GAG CCG CTT TTT G, rev: AAT TCA AAA AGC GGC TCT CTC TAC TAC TTC ATG GGT CAG GTG AAG TAG TAG AGA GAG CCG C); *shFezf2#2* (fwd: CTA GGG TGT TCA ATG CTC ACT ATA ACC TGA CCC ATT ATA GTG AGC ATT GAA CAC CTT TTT G, rev: AAT TCA AAA AGG TGT TCA ATG CTC ACT ATA ATG GGT CAG GTT ATA GTG AGC ATT GAA CAC C); *shFezf2#3* (fwd: CTA GGC TGC TCA GTT ACT CGG AGT TCC TGA CCC AAA CTC CGA GTA ACT GAG CAG CTT TTT G, rev: AAT TCA AAA AGC TGC TCA GTT ACT CGG AGT TTG GGT CAG GAA CTC CGA GTA ACT GAG CAG C), *shFezf2#4* (fwd: CTA GGC TAG ACC GTT TGT GTG CAA ACC TGA CCC ATT TGC ACA CAA ACG GTC TAG CTT TTT G, rev: AAT TCA AAA AGC TAG ACC GTT TGT GTG CAA ATG GGT CAG GTT TGC ACA CAA ACG GTC TAG C); *shFezf2#5* (fwd: CTA GGA GTC AAG AGC CAC AGC AAA CCC TGA CCC AGT TTG CTG TGG CTC TTG ACT CTT TTT G, rev: AAT TCA AAA AGA GTC AAG AGC CAC AGC AAA CTG GGT CAG GGT TTG CTG TGG CTC TTG ACT C); *shCdh22#6* (fwd: CTA GGC AGA CGA CCC TAC TTA CAC GCC TGA CCC ACG TGT AAG TAG GGT CGT CTG CTT TTT G, rev: AAT TCA AAA AGC AGA CGA CCC TAC TTA CAC GTG GGT CAG GCG TGT AAG TAG GGT CGT CTG C); *shCdh22#7* (fwd: CTA GGG ACA ATG TGA TCA AAT AAC GCC TGA CCC ACG TTA TTT GAT CAC ATT GTC CTT TTT G, rev: AAT TCA AAA AGG ACA ATG TGA TCA AAT AAC GTG GGT CAG GCG TTA TTT GAT CAC ATT GTC C); *shCdh22#8* (fwd: CTA GGG CTG GCA CAA CAT TAC TAC GCC TGA CCC ACG TAG TAA TGT TGT GCC AGC CTT TTT G, rev: AAT TCA AAA AGG CTG GCA CAA CAT TAC TAC GTG GGT CAG GCG TAG TAA TGT TGT GCC AGC C); *shCntn6#6* (fwd: CTA GGG AGC ACA TCT CTA GTA CAC GCC TGA CCC ACG TGT ACT AGA GAT GTG CTC CTT TTT G, rev: AAT TCA AAA AGG AGC ACA TCT CTA GTA CAC GTG GGT CAG GCG TGT ACT AGA GAT GTG CTC C); *shCntn6#7* (fwd: CTA GGA AAT CCA AAT CCT TCG TAC GCC TGA CCC ACG TAC GAA GGA TTT GGA TTT CTT TTT G, rev: AAT TCA AAA AGA AAT CCA AAT CCT TCG TAC GTG GGT CAG GCG TAC GAA GGA TTT GGA TTT C). We used a *shLacZ* sequence as control, and the shRNAs were cloned in a backbone plasmid as described before^11^.

### In vitro validation of shRNA sequences

To assess the *in vitro* efficacy of *shRNA* by Western Blot, Cre-expressing Human embryonic kidney (HEK, Ambion #SC004-BSD) cells were transfected with plasmids expressing cDNA and either *shLacZ* or the *shRNA* against the target gene. 48-72 hours post-transfection, HEK293T were rinsed with 1x ice-cold PBS. Samples were homogenized in lysis buffer containing 25 mM Tris-HCl pH 8, 50mM NaCl, 1% Triton X-100, 0.5% sodium deoxycholate, 0.001 % SDS and protease inhibitor cocktail (cOmplete Mini, Roche). Three replicates were assessed for each *shRNA* sequence tested. Samples were resolved by SDS-PAGE and transferred onto PVDF membranes. Membranes were blocked with blocking solution consisting of 5% Blotting-Grade Blocker (Bio-Rad, #1706404) in TBST (20mM Tris-HCl pH7.5, 150mM NaCl and 0.1% Tween20) for 1 hour and incubated with mouse anti-HA (Invitrogen #26183, 1:500) in blocking solution overnight at 4°C. After washing 3 x 10 min in TBST, membranes were incubated with Goat anti-mouse HRP (Invitrogen #31444, 1:5,000) in blocking solution for 1 hour at RT, followed by 3 x 10 min washes in TBST. Protein levels were visualized by chemiluminescence using either Immobilon Chemiluminescent HRP substrate ECL (Millipore #WBKLS0500) or SuperSignal West Femto Maximum Sensitivity Substrate (Thermo Fisher #34095), depending on signal intensity. Blots were visualized using a LI-COR Odyssey Fc imaging system. Then, blots were washed in TBST and incubated with mouse anti-ß-actin HRP (Sigma-Aldrich # A3854, 1:20,000) in blocking solution for 1 hour at RT. Actin signal was visualized using SuperSignal West Pico Chemiluminescent substrate ECL (Thermo Fisher #34577). For each sample, HA signal intensity was normalized to the actin signal. The resulting value for each sample was then normalized to the *shLacZ* value within the same replicate. For *Fezf2*, *shRNA* #1 and #2 were chosen. *shRNA* virus was made by co-transfecting both plasmids. For *Cdh22* and *Cntn6*, robust downregulation was only observed for one sequence. Therefore, only that sequence (*Cdh22#7* and *Cntn6#6*) was selected for viral production.

### Virus production

For the in-house production of AAVs, HEK293FT cells (Thermo Fisher Scientific) were cultured in Dulbecco’s Modified Eagle’s medium supplemented with 10% fetal bovine serum, 2 mM glutamine, penicillin (50 units/ml), streptomycin (50 g/ml) 200 mM Hepes (Thermo Fisher Scientific) and Gibco MEM Non-Essential Amino Acids. The cultures were incubated at 37°C in a humidified atmosphere containing 5% CO2. HEK293T cells were transfected using polyethylenimine (PEI, Sigma) at a 1:4 DNA:PEI ratio or Lipofectamine 2000 (Thermo Fisher Scientific). HEK293FT cells were seeded on 15-cm plates and co-transfected with packaging plasmids AAV-ITR-2 genomic vectors (7.5 µg), AAV-Cap8 vector pDP8 (30 µg; PlasmidFactory GmbH #pF478) or AAV-Cap DJ Rep-Cap vector (30 µg; Cell Biolabs #VPK-420-DJ) using PEI (Sigma) at a ratio 1:4 (DNA:PEI). 72 hours post-transfection, supernatants were incubated with Ammonium sulfate (65g/200ml supernatant) for 30 minutes on ice and centrifuged for 45 minutes at 4000 RPM at 4°C. Transfected cells were harvested and lysed (150mM NaCl, 50mM Tris pH8.5), followed by three freeze-thaw cycles and Benzonase treatment (50U/ml; Sigma) for 1 hour at 37°C. Filtered AAVs (0.8 µm and 0.45 µm MCE filters) from supernatants and lysates were run on an Iodixanol gradient by ultracentrifugation (Vti50 rotor, Beckmann Coultier) at 50,000 RPM for 1 hour at 12°C. The 40% iodixanol fraction containing the AAVs was collected, concentrated using 100-kDa molecular weight cut-off (MWCO) filters Centricon plus-20 and Centricon plus-2 (Merck-Millipore), aliquoted and stored at - 80°C. The number of genomic copies was determined by qPCR using the following primers against the WPRE (Fwd: GGC ACT GAC AAT TCC GTG GT and Rev: CGC TGG ATT GAG GGC CGAA).

Home-made viruses used in this study were: *hSyn-DIO-tdTomato* (3.58 x 10^13^ vg/ml), *hSyn-DIO-Kir2.1-T2A-tdTomato* (2.63 x 10^13^ vg/ml), *hSyn-DIO-mutKir2.1-T2A-tdTomato* (4.69 x 10^13^ vg/ml), *hSyn-DIO-HA-Fezf2-T2A-EGFP* (9.72 x 10^11^ vg/ml), *EF1a-DIO-EGFP-shLacZ* (8.00 x 10^11^ vg/ml), *EF1a-DIO-EGFP-shFezf2* (2.53 x 10^12^ vg/ml), *EF1a-DIO-EGFP-shCdh22* (9.07 x 10^12^ vg/ml), *EF1a-DIO-EGFP-shCntn6* (1.30 x 10^13^ vg/ml).

### Commercial viruses

The following commercial viruses were produced by AddGene: *rAAVretro-EF1a-Cre* (2.1 x 10^13^ vg/ml, #55636-AAVrg, a gift from Karl Deisseroth), *AAV8-hSyn-DIO-hM3D(Gq)-mCherry* (1.8 x 10^13^ vg/ml, #44361-AAV8, a gift from Bryan Roth), and *AAV8-hSyn-DIO-GFP* (2.3 x 10^13^ vg/ml, #50457-AAV8, a gift from Bryan Roth), *AAV8-hSyn-DIO-EGFP* (2.3 x 10^13^ vg/ml, #50457-AAV8, a gift from Bryan Roth). *rAAVretro-hSyn-CreERT2* was obtained from the Viral Vector Facility of the University of Zürich (7.9 x 10^12^ vg/ml, #v549).

### Viral injections

P1 pups were anaesthetized with isoflurane and mounted on a stereotactic frame (Stoelting). To target ET pyramidal neurons in wild-type animals, we used an intersectional strategy. *rAAV2retro-EF1a-Cre* was injected in the pontine nuclei (AP -1.2 mm, ML 0.3, DV 4.2 and 4.0 mm from lambda; 150 nl per depth). For experiments involving inducible Cre, *rAAVretro-hSyn-CreERT2* was injected instead. In the same animals, relevant Cre-dependent AAVs were injected in the ipsilateral S1 at a depth of 0.3 mm (single injection, 300 nl). For *Fezf2* overexpression experiments, AAVs were injected at P3 to prevent any possible identity change. For experiments using Cre lines, the virus was only injected into the S1. Injection speed was 200 nl/min for all injections.

### *In vitro* electrophysiology

For all experiments, an experimental animal and an age-matched – and where possible sex-matched – littermate control was used for electrophysiology on the same day. Mice were deeply anesthetized with an overdose of sodium pentobarbital and transcardially perfused with 10 mL ice-cold slicing solution containing (in mM): 87 NaCl, 75 sucrose, 26 NaHCO3, 11 glucose, 7 MgCl2, 2.5 KCl, 1.25 NaH2PO4 and 0.5 CaCl2, oxygenated with 95% O2 and 5% CO2. The brain was quickly removed, and the injected hemisphere was glued to a cutting platform before being submerged in ice-cold slicing solution. 300 µm thick coronal slices containing S1 were cut using a vibratome (Leica VT1200S, Wetzlar, Germany) and incubated for 45-60 min at 32°C, and subsequently at room temperature, in the same solution. All salts were purchased from Sigma-Aldrich (St. Louis, MO). Slices were transferred to the recording setup and superfused with recording ACSF containing (in mM) 124 NaCl, 1.25 NaH2PO4, 3 KCl, 26 NaHCO3, 10 Glucose, 2 CaCl2, and 1 MgCl2, which was oxygenated with 95% O2 and 5% CO2 and heated to 34°C. Pipettes (3–5 MΩ) were made from borosilicate glass capillaries using a PC-10 pipette puller (P10, Narishige, London, UK) and filled with intracellular solution containing (in mM) 70 K-gluconate, 70 KCl, 10 Hepes, 4 Mg-ATP, 4 K2-phosphocreatine, and 0.4 GTP, adjusted with KOH to pH 7.3 (±290 mOsm). Where applicable, 1% neurobiotin (Vectorlabs) was added to the intracellular solution.

Recordings were made using a Multiclamp 700B amplifier (Molecular Devices, San Jose, CA). The signal was passed through a Hum Bug Noise Eliminator (Digitimer, Welwyn Garden City, UK), sampled at 10 kHz, and filtered at 3 kHz using a Digidata 1440A (Molecular Devices, San Jose, CA). Cells were excluded if the access resistance (Ra) exceeded 30 MΩ. Resting membrane potential was measured in current-clamp for 5 – 10 s immediately after breaking in. Intrinsic properties were measured by applying a series of negative or positive current injections while maintaining a baseline Vm of -70 mV, at a step size of 25 pA. Active and passive properties were analysed in Matlab (Mathworks, Natick, MA) using custom scripts. Input resistance was calculated as the linear slope of the current-voltage (I-V) relationship of the last 200 ms of all negative stimuli. Voltage sag was measured during the current step in which the peak voltage deflection was nearest -90 mV, and was calculated as the percentage change between the peak of the response and the average voltage deflection of the last 200 ms of the same step. Rheobase was determined as the average first current injection to elicit and action potential across three series of current steps at a step size of 2 pA. Miniature inhibitory postsynaptic currents (mIPSCs) recordings were performed at a holding voltage of -76 mV in the presence of 1 µM tetrodotoxin (TTX, HB1035) 10 µM 2,3-Dioxo-6-nitro-1,2,3,4-tetrahydrobenzo[f]quinoxaline-7-sulfonamide (NBQX, HB0443) and 50 µM D-(−)-2-Amino-5-phosphonopentanoic acid (D-APV, HB0225), all of which were purchased from Hello Bio (Bristol, UK). mIPSCs were analyzed using MiniAnalysis (SynaptoSoft, Decatur, GA, USA).

### RNA sequencing

#### Ribosome-associated RNA isolation

Mice were deeply anesthetized with an overdose of sodium pentobarbital. The infected cortex was rapidly dissected in ice-cold RNase-free PBS. Lower layers of S1 were dissected out and immediately homogenised in ice-cold lysis buffer (50 mM Tris-HCl pH 7.5, 100 mM KCl, 12 mM MgCl2, 1 mg/ml Heparin (Sigma-Aldrich), cOmplete EDTA-free protease inhibitors (Sigma-Aldrich), 200 U/ml RNAsin (Promega), 100 µg/ml cycloheximide and 1 mM DTT (Sigma-Aldrich)). Tissue from 2-3 brains were pooled per sample. Samples were centrifuged at 2,000 g for 10 min at 4°C, and Igepal-CA630 (Sigma-Aldrich) was added to the samples to a final concentration of 1%. Samples were then centrifuged at 13,000 g for 10 min at 4°C. The supernatant was added on 100 µl of anti-HA magnetic beads (Pierce #88837) for 3-4 h at 4°C with gentle rotation. After incubation, beads were washed 3 times in ice-cold washing buffer (300 mM KCl, 1% Igepal-CA630, 50 mM Tris-HCl pH 7.5, 12mM MgCl2, 1mM DTT and 100 µg/ml cycloheximide) and eluted in 350 µl of RLT Plus buffer (RNAeasy Plus Micro kit, Qiagen) supplemented with 2-mercaptoethanol (Bio-Rad).

#### RNA purification, library preparation and sequencing

RNA purification of immunoprecipitated RNA was performed using the RNeasy Plus Micro kit (Qiagen) following the manufacturer’s protocol. The quality of immunoprecipitated RNA was verified on a Bioanalyzer instrument (Agilent Technologies) using an RNA 6000 Pico Chip. All RNA samples had RNA integrity number (RIN) values of 9.8 or higher. Three biological replicates were analyzed for each genotype. Library preparation and RNA-seq experiments were performed by the Genomic Unit of the Centre for Genomic Regulation (CRG, Barcelona, Spain). Libraries were prepared using NEBNext Poly(A) mRNA Magnetic Isolation Module (New England Biolabs, #7490) and NEBNext Ultra II Directional RNA Library Prep Kit for Illumina (New England Biolabs, 7760) according to the manufacturer’s protocol. Libraries were then sequenced paired-end using an Illumina HiSeq 2500 platform to a minimum of 100 million mapped reads per sample.

#### RNA sequencing analysis

Bioinformatic analyses were carried out by GenoSplice (Paris, France). Briefly, Data quality was assessed using FastQC and rseqc^71^. Reads were mapped using STAR_2.4.0^72^ to reference genome mm10. Counts were obtained through FAST DB (v2018_1), gene annotations (GenoSplice) and normalization using Deseq. One sample (mutKir1) did not cluster sufficiently and was discarded from further differential gene analysis and selection of target genes. Gene Ontology (GO) analysis was performed using WebGestalt (https://academic.oup.com/nar/article/47/W1/W199/5494758) using the terms biological process, cellular component, and molecular function, as well as the KEGG and REACTOME pathway databases. GO terms were considered to be significantly enriched at FDR < 0.05.

To select gene targets, differentially expressed genes within the following Panther GO terms were considered: *cell adhesion molecule* (PC00069), *scaffold/adaptor protein* (PC00226) and *C2H2 zinc finger transcription factor* (PC00248). Genes were then ranked using three criteria. First, genes were scored according to the expression fold-change between Kir2.1 and mutKir2.1. Second, because low-count genes are often unreliable, genes were ranked according to their expression levels in mutKir2.1 controls. Third, to bias towards ET-specific genes, genes were scored according to enrichment in ET cells vs IT cells using single-cell sequencing data from the Allen Brain Institute database^73^. These criteria were converted into a single rank score by calculating Z-scores for each criterion and averaging the absolute Z-score of criterion 1 and the Z-scores of criteria 2 and 3. Rankings for all criteria and the final rank score are shown in Figure 2D.

### Histology

#### Immunohistochemistry

Animals were deeply anaesthetized with sodium pentobarbital by intraperitoneal injection and transcardially perfused with 0.9% saline followed by 4% paraformaldehyde (PFA) in PBS. Dissected brains were post-fixed for 2 h at 4°C in 4% PFA and cryoprotected in successive solutions of 15% and 30% sucrose (Sigma-Aldrich, Cat# S0389) in PBS. 40 µm thick sections containing S1 were cut using a sliding microtome (Leica SM2010 R) and stored in ethylene glycol (30% ethylene glycol (Merck, 324558), 30% Glycerol (Merck, G5516) in PBS) at -20 °C. Free-floating sections were permeabilized with 0.25% Triton X-100 (Sigma-Aldrich, Cat# T8787) in PBS for 4 times 15 min at room temperature (20-22 °C, RT) and incubated for 2 hours at RT in a solution containing 0.3% Triton X-100, 5% bovine serum albumin (BSA) (Sigma-Aldrich, Cat# A8806) and 10% normal goat serum and/or normal donkey serum, corresponding to the secondary antibodies used. Sections were then incubated overnight at 4 °C with primary antibodies in 0.3% Triton X-100, 2% BSA and 5% normal serum. Sections were washed 3 times 10 min in PBS and incubated with secondary antibodies in 0.3% Triton X-100, 2% BSA and 5% serum for 2 hours at RT. The following primary and secondary antibodies were used: mouse anti-NeuN (Sigma-Aldrich #MAB377, 1:500), rabbit anti-NeuN (Millipore # ABN78, 1:500), goat anti-mCherry (Antibodies-Online #ABIN1440057, 1:500), rabbit anti-DsRed (Clontech #632496, 1:500), chicken anti-GFP (Aves Lab #GFP-1020, 1:1000), Mouse IgG2a anti-Syt2 (ZFIN #ZDB-ATB-081002-25, 1:250), Mouse IgG1 anti-HA (BioLegend #901502, 1:330), Rabbit anti-CFOS (SySy #AbE 457, 1:500). Goat anti-mouse IgG1 488 (Invitrogen #A-21121, 1:500), Goat anti-chicken 488 (Invitrogen #A-11039, 1:500), Donkey anti-rabbit 488 (Invitrogen #A-21206, 1:500), Donkey anti-rabbit 555 (Invitrogen #A-31572, 1:500), Donkey anti-goat 555 (Invitrogen #A-21432, 1:500), Goat anti-mouse IgG1 555 (Invitrogen #A-21127, 1:500), Goat anti-mouse IgG2a 647 (Invitrogen #A-21241, 1:500).

For the physiological validation of the function of *AAV-hSyn-DIO-Kir2.1-RPL10a*, cells were filled with neurobiotin during recording, and slices were subsequently fixed in PFA for 1 hour at RT. Slices were incubated with 1:500 streptavidin-488 (Thermo Fisher) in PBS containing 0.25% triton for 24 hours at 4 °C. Immunohistochemistry was then performed as described above using mouse IgG1 anti-HA and donkey anti-mouse IgG1-555.

#### Single-molecule fluorescence in situ hybridization

Animals were perfused as previously described. Brains were dissected and postfixed overnight in 4% PFA at 4 °C, followed by cryoprotection in 15% and 30% sucrose in RNase-free PBS at 4 °C. 30 µm thick sections were cut and stored in RNase-free ethylene glycol solution at -70 °C. Sections were mounted on RNase-free SuperFrost Plus slides (Epredia # J1800AMNZ) and dehydrated using successive steps of 50%, 70% and 100% ethanol.

Single-molecule fluorescence *in situ* hybridization (smFISH) was performed using the RNAscope Multiplex Fluorescent Assay v2 protocol (ACDBio #323110) according to the manufacturer’s instructions. *Fezf2* (ACDBio #313301 and ACDBio #313301-C2), *Cdh22* (ACDBio #573541-C3) and *Cntn6* (ACDBio #836461) mRNA was visualized using Opal 570 (Akoya BioScience, FP1488001KT) and Opal 650 (Akoya BioScience, FP1496001KT).

Immunohistochemistry was then performed to visualize infected neurons. Slide-mounted sections were incubated in blocking buffer containing 10% normal goat serum for 30 min at RT, and incubated with primary antibodies Chicken anti-GFP (Aves Lab #GFP-1020, 1:500) and Chicken anti-GFP (Abcam #ab13970, 1:500) overnight at 4 °C. After washing, sections were incubated with Goat anti-chicken 488 (Molecular Probes #A-11039) for 2 hours at RT.

### Image acquisition and analysis

For analysis of perisomatic boutons, z-stacks were acquired on an inverted Leica TCS-SP8 confocal using a 100X numerical aperture (NA) 1.44 oil immersion objective at a voxel size (xyz) of 0.05 x 0.05 x 0.2 µm maintaining the same laser power, photomultiplier gain, pinhole, and detection filter settings for all animals within an experiment and analyzed with IMARIS 7.5.2 software. Somata were manually reconstructed from NeuN signal using the Surface function. The detection threshold for Syt2 puncta was adjusted for staining intensity per animal and Syt2 puncta were automatically detected using the Spots function. The number of Syt2 puncta close to the reconstructed soma was quantified using the “Find spots close to surface” tool (ImarisXT extension) with a distance threshold of 0.4 µm.

Low-magnification confocal images were obtained using a 10x NA 0.3 dry objective. c-Fos-expressing cells were quantified using the Cell Counter plugin in FIJI (Schindelin et al., 2012).

For analysis of smFISH, single plane images were acquired on an inverted Leica TCS-SP8 confocal using a 63X NA 1.4 oil immersion objective at a pixel size (xy) of 0.09 x 0.09 µm. Images were analyzed using custom Matlab (Mathworks) scripts. The regions of interest (ROIs) covering the soma of infected cells were manually delineated using the fluorescent reporter signal, and mRNA signal was measured for each infected cell as the surface of the mRNA signal within the ROI divided by the total surface of the ROI in percentages. Because RNAscope signal quality can vary significantly between animals and even within sections, for each image the value of each ROI was normalized to the brightest uninfected cell in the image. For both quantification of synaptic boutons and smFISH, a minimum of 15 cells were analyzed per animal, and the average value per animal was used for statistical analysis.

To determine whether recorded and neurobiotin-filled cells were positive for HA, cells were imaged using a Zeiss ApoTome with a 63x NA 1.4 objective. Only cells with strong HA signal were included in subsequent analysis of electrophysiological recordings.

### Statistical analyses

All statistical tests were performed in Prism 10 (Graphpad). Parametric data were analyzed using a Student’s t-test in the case of equal variances between groups and a Welch’s t-test in the case of unequal variance (>2-fold difference in standard deviation between groups). For *in vitro* validation of *shRNA* sequences, each sequence was compared to *shLacZ* using a t-test and Bonferroni correction was applied. For analysis of the correlation of mRNA signal values in the *Fezf2* overexpression experiments, a simple linear regression was used. P values < 0.05 were considered statistically significant. All data are presented as mean ± SEM.

